# Multiparametric Platform for Profiling Lipid Trafficking in Human Leukocytes: Application for Hypercholesterolemia

**DOI:** 10.1101/2021.04.19.440471

**Authors:** Simon G. Pfisterer, Ivonne Brock, Kristiina Kanerva, Iryna Hlushchenko, Lassi Paavolainen, Pietari Ripatti, Mohammad M. Islam, Aija Kyttälä, Maria D. Di Taranto, Annalisa Scotto di Frega, Giuliana Fortunato, Johanna Kuusisto, Peter Horvath, Samuli Ripatti, Markku Laakso, Elina Ikonen

## Abstract

Systematic insight into cellular dysfunctions can improve understanding of disease etiology, risk assessment and patient stratification. We present a multiparametric high-content imaging platform enabling quantification of low-density lipoprotein (LDL) uptake and lipid storage in cytoplasmic droplets of primary leukocyte subpopulations. We validated this platform with samples from 65 individuals with variable blood LDL-cholesterol (LDL-c) levels, including familial hypercholesterolemia (FH) and non-FH subjects. We integrated lipid storage data into a novel readout, lipid mobilization, measuring the efficiency with which cells deplete lipid reservoirs. Lipid mobilization correlated positively with LDL uptake and negatively with hypercholesterolemia and age, improving differentiation of individuals with normal and elevated LDL-c. Moreover, combination of cell-based readouts with a polygenic risk score for LDL-c explained hypercholesterolemia better than the genetic risk score alone. This platform provides functional insights into cellular lipid trafficking from a few ml’s of blood and is applicable to dissect metabolic disorders, such as hypercholesterolemia.

**Motivation:** We have limited information on how cellular lipid uptake and processing differ between individuals and influence the development of metabolic diseases, such as hypercholesterolemia. Available assays are labor intensive, require skilled personnel and are difficult to scale to higher throughput, making it challenging to obtain systematic functional cell-based data from individuals. To overcome this problem, we established a scalable automated analysis pipeline enabling reliable quantification of multiple cellular readouts, including lipid uptake, storage and mobilization, from different white blood cell populations. This approach provides new personalized insights into the cellular basis of hypercholesterolemia and obesity.

**Graphical Abstract:** 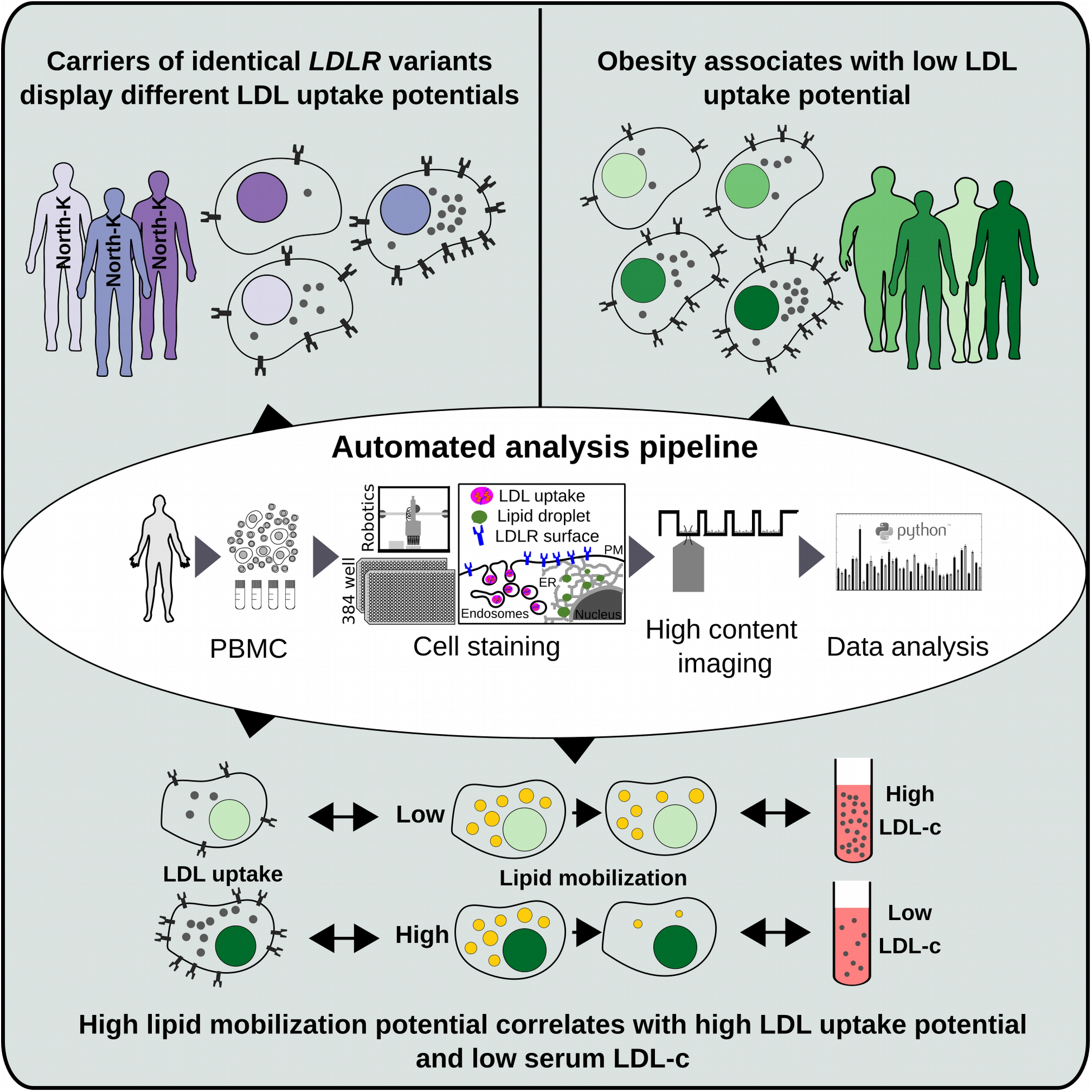

## Introduction

Hypercholesterolemia is one of the most common metabolic disorders and a major risk factor for cardiovascular disease (CVD). It is characterized by an accumulation of low-density lipoprotein cholesterol (LDL-c) in the blood^1^. In familial hypercholesterolemia (FH), mutations, most commonly in the LDL receptor (*LDLR*) gene, lead to increased LDL-c. However, FH represents only 2.5% of all hypercholesterolemia patients. For the remainder, polygenic and lifestyle effects appear as the main contributing factors^2–5^.

So far, we have little information on how cellular lipid trafficking and storage are altered in individual patients. However, systematic assessment of LDL uptake and other mechanisms related to hypercholesterolemia could provide insights into disease mechanisms and treatment outcomes in a personalized manner. The majority of high-risk hypercholesterolemia patients does not achieve their LDL-c target levels^6^. This could be due to sub-optimal treatment, non-adherence to therapy and/or cellular programs limiting drug efficacy. Increased evidence from cancer therapy demonstrates that cell-based assays can provide better targeted and more effective personalized treatment strategies^7^. Regarding hypercholesterolemia, we need to establish scalable and reliable assays that allow systematic profiling of functional defects in individual persons and evaluate how to utilize such assays to better explain factors contributing to hypercholesterolemia in individual patients.

The currently used cell-based assays for studying the etiology of hypercholesterolemia are quantification of cellular LDL uptake or LDLR cell surface expression using flow cytometry. These readouts have been mostly utilized to characterize the severity of *LDLR* mutations in FH patients^8,9^. However, LDLR surface expression and LDL uptake are highly variable among FH patients^10–12^. This not only speaks for the importance of functional cell-based assays but also calls for new cellular readouts to better characterize the heterogeneity of lipid metabolism in individual subjects.

LDLR expression and cellular LDL internalization are tightly regulated. Low cholesterol levels in the endoplasmic reticulum (ER) signal cholesterol starvation and trigger increased LDLR expression, while high cholesterol in the ER downregulates LDLR expression. Excess ER cholesterol is stored as cholesterol ester in lipid droplets (LD), from where it can be mobilized upon need^13,14^. We therefore considered that quantification of cellular LDs and their dynamic changes upon altering lipoprotein availability may provide additional information for assessing the cellular basis of hypercholesterolemia.

Here, we established sensitive and scalable analyses for automated quantification of fluorescent lipid uptake, storage and removal in primary lymphocyte and monocyte populations, and defined lipid mobilization as a novel parameter measuring how efficiently cells deplete their lipid stores. We found marked differences in the parameters established in both FH and non-FH study groups and highlight their potential to provide deeper insights into the cellular mechanisms of hypercholesterolemia.

## Results

### Automated pipeline for quantification of hypercholesterolemia-related functional defects in primary human leukocytes

Several cell types such as lymphocytes, monocytes and Epstein-Barr virus (EBV) immortalized lymphoblasts have been used for measuring LDL uptake^15,16^. Whilst EBV lymphoblasts show the highest LDL uptake, cell immortalization is time consuming and alters cellular functions^15,17^. We therefore set up an automated imaging and analysis pipeline for sensitive quantification of LDL uptake and LDLR surface expression from less than two million peripheral blood mononuclear cells (PBMCs) (**Figure 1A**). Cryopreserved PBMCs were recovered in 96-well plates at defined densities and incubated with lipid-rich control medium (CM, 10% FBS) or lipid poor medium (LP, 5% lipoprotein-deficient serum) for 24 h. Cells were labeled with fluorescent LDL particles (DiI-LDL) for 1 h, washed and automatically transferred to 384-well plates for staining and automated high-content imaging (**Figure 1A**). After adhesion to coated imaging plates, lymphocytes remain small while monocytes spread out, enabling a crude classification of leukocyte populations based on size: PBMCs with a cytoplasmic area <115 μm^2^ were classified as a lymphocyte-enriched fraction (from here on lymphocytes) and those with a cytoplasmic area >115 μm^2^ as monocyte-enriched fraction (from here on monocytes) (**Suppl. Figure 1A-C**).

**Figure 1:**
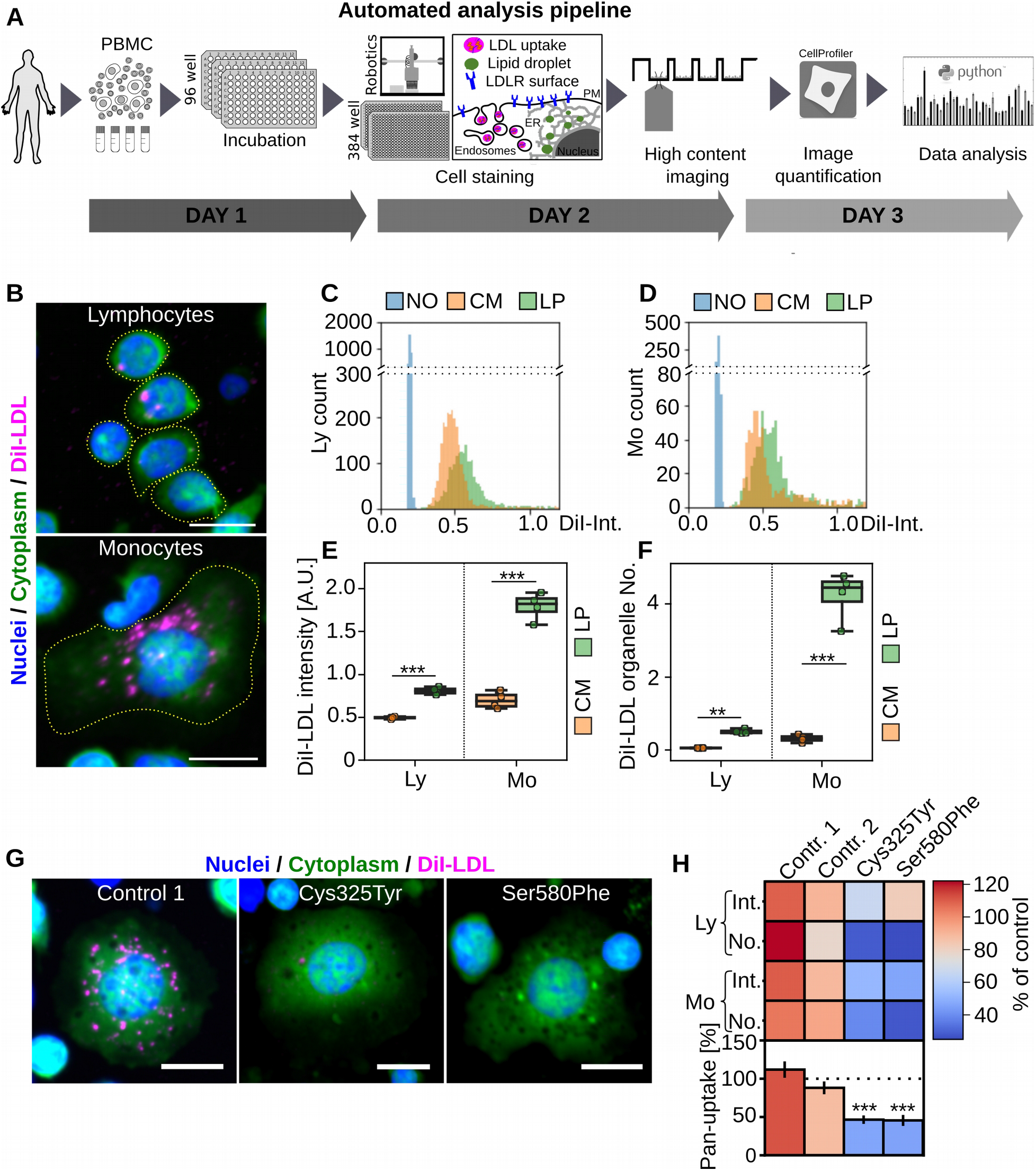
Automated analysis pipeline for multiplex quantification of functional phenotypes in PBMCs. **A**) Schematic presentation of the automated analysis pipeline. For each experiment cryopreserved PBMC samples were thawed, aliquoted into 96 wells and incubated overnight with lipid rich (CM, 10% FBS) or lipid poor (LP, 5% LPDS) medium. Cells were labeled with fluorescent LDL (DiI-LDL) or directly transferred to 384 well imaging plates, automatically fixed, stained and subjected to automated high-content imaging. Images were quantified with CellProfiler and single-cell data was processed with Python tools. **B**) Representative images of lymphocyte and monocyte DiI-LDL uptake after lipid starvation. **C**) Histogram for cellular DiI-LDL intensities in lymphocytes and monocytes (**D**) from a single well. **E**) Quantification of mean DiI-LDL intensities and DiI-LDL organelles (**F**) in lymphocytes (Ly) and monocytes (Mo); representative of eight independent experiments, each with four wells per treatment; Student’s t-test. **G**) Representative images of DiI-LDL uptake in monocytes isolated from FH patients with LDLR mutations Cys325Tyr or Ser580Phe and a control after lipid starvation. **H**) Quantification of monocyte (Mo) and lymphocyte (Ly) cellular DiI-LDL intensities (Int), DiI-LDL organelle numbers (No) and pan-uptake; duplicate wells / patient (eight wells / patient for pan-uptake). Significant changes to control 2 were calculated with Welch’s t-test. ***p < 0.001, **p < 0.01, scale bar = 10 μm, error bars = SEM.

In CM, DiI-LDL uptake into lymphocytes and monocytes was more than two-fold above the background of non-labeled cells (**Figure 1B-D**). Lipid starvation further increased DiI-LDL uptake in both cell populations, as expected (**Figure 1C, D**). We quantified about 700 monocytes and 2300 lymphocytes per well (**Suppl. Figure 1D**), aggregated the single-cell data from individual wells and averaged the results from 2-4 wells for each treatment (**Suppl. Figure 1D**). For both cell populations, we defined two readouts, cellular DiI-LDL intensity (DiI-Int), reflecting DiI-LDL surface binding and internalization, and DiI-LDL organelle number (DiI-No), reflecting internalized DiI-LDL (**Figure 1E, F**). This resulted in four parameters: Monocyte (Mo) DiI-Int, lymphocyte (Ly) DiI-Int, Mo DiI-No and Ly DiI-No. In both cell populations, DiI-Int was inhibited by adding surplus unlabeled LDL, arguing for a saturable, receptor-mediated uptake mechanism (**Suppl. Figure 1E**).

In lipid rich conditions, Mo DiI-Int was slightly higher than Ly DiI-Int (**Figure 1E**), and upon lipid starvation, Mo DiI-Int increased more substantially, providing a larger fold increase than Ly DiI-Int (**Figure 1E**). Furthermore, Mo DiI-No was roughly ten-fold higher than Ly DiI-No, with both parameters showing a five-fold increase upon lipid starvation (**Figure 1F**). Thus, DiI-LDL uptake into monocytes was better than into lymphocytes, but both cell populations responded to lipid starvation. As EBV lymphoblasts are often a preferred choice for LDL uptake studies^20^ we compared LDL uptake between EBV lymphoblasts and monocytes (**Suppl. Figure 1F,G**). This showed that DiI-Int signal after lipid starvation was roughly similar in EBV-lymphoblasts and monocytes, implying that the primary cells provide high enough DiI-LDL signal intensities without cell immortalization (**Suppl. Figure 1G**).

To enable data comparison between experiments, we included two controls. Each control consisted of a mixture of large-scale PBMC isolations from four healthy blood donors, with the cells cryopreserved at a defined density for one-time use aliquots. In each experiment, Mo DiI-Int, Ly DiI-Int, Mo DiI-No and Ly DiI-No were normalized to these controls. We also introduced a combinatorial score, pan-LDL uptake (or pan-uptake), representing the average of Mo DiI-Int, Ly DiI-Int, Mo DiI-No and Ly DiI-No. We then assessed the intraindividual variability of these five readouts in three individuals on two consecutive days (**Suppl. Figure 1H**). The intraindividual variability was low for a cell-based assay, especially in monocytes, with 7.6% for Mo DiI-No, 12% for Mo DiI-Int and 13% for pan-uptake. The values were only moderately higher in lymphocytes, with DiI-Int 15% and DiI-No 21% variability (**Suppl. Figure 1I**).

We next validated our LDL uptake measurements in PBMCs of two He-FH patients with highly elevated LDL-c and reduced LDL uptake in EBV lymphoblasts (Cys325Tyr and Ser580Phe mutations in *LDLR*) (**Suppl. Figure 1J**). For both patients, Mo and Ly DiI-No as well as Mo DiI-Int were reduced by more than 45%, Ly DiI-Int was less profoundly decreased, and pan-uptake was reduced by over 50% (**Figure 1G, H; Suppl. Figure 1J**). Together, these data indicate that our analysis pipeline enables quantification of multiple LDL uptake parameters in major leukocyte cell populations and distinguishes defective LDLR function therein.

### Heterogeneous LDL uptake and LDLR surface expression in He-FH patients

We next used this pipeline to characterize 21 He-FH patients from the Metabolic Syndrome in Men (METSIM) cohort study^18^ (**Suppl. Table 1**). The patients’ mutations reside in the *LDLR* coding region and range from pathogenic to likely benign variants (**Figure 2A**). Quantification of DiI-Int and DiI-No for monocytes and lymphocytes provided relatively similar results for each individual (**Figure 2B**). However, there were substantial differences in these parameters between individuals, including patients harboring identical *LDLR* mutations (**Figure 2B**). This was most pronounced for FH-North Karelia (Pro309Lysfs*59), a pathogenic loss-of-function variant, but also evident for FH-Pogosta (Arg595Gln) and FH-Glu626Lys (**Figure 2A, B**). These observations imply that in He-FH, regulatory mechanisms may enhance the expression of the unaffected *LDLR* allele and/or stabilize the encoded protein. In support of this notion, we obtained a strong correlation between monocyte LDLR surface expression and DiI-Int, DiI-No and pan-uptake scores for the same individuals (pan-uptake, R=0.58, p=0.006), (**Figure 2C, Suppl. Figure 2A**).

**Figure 2).**
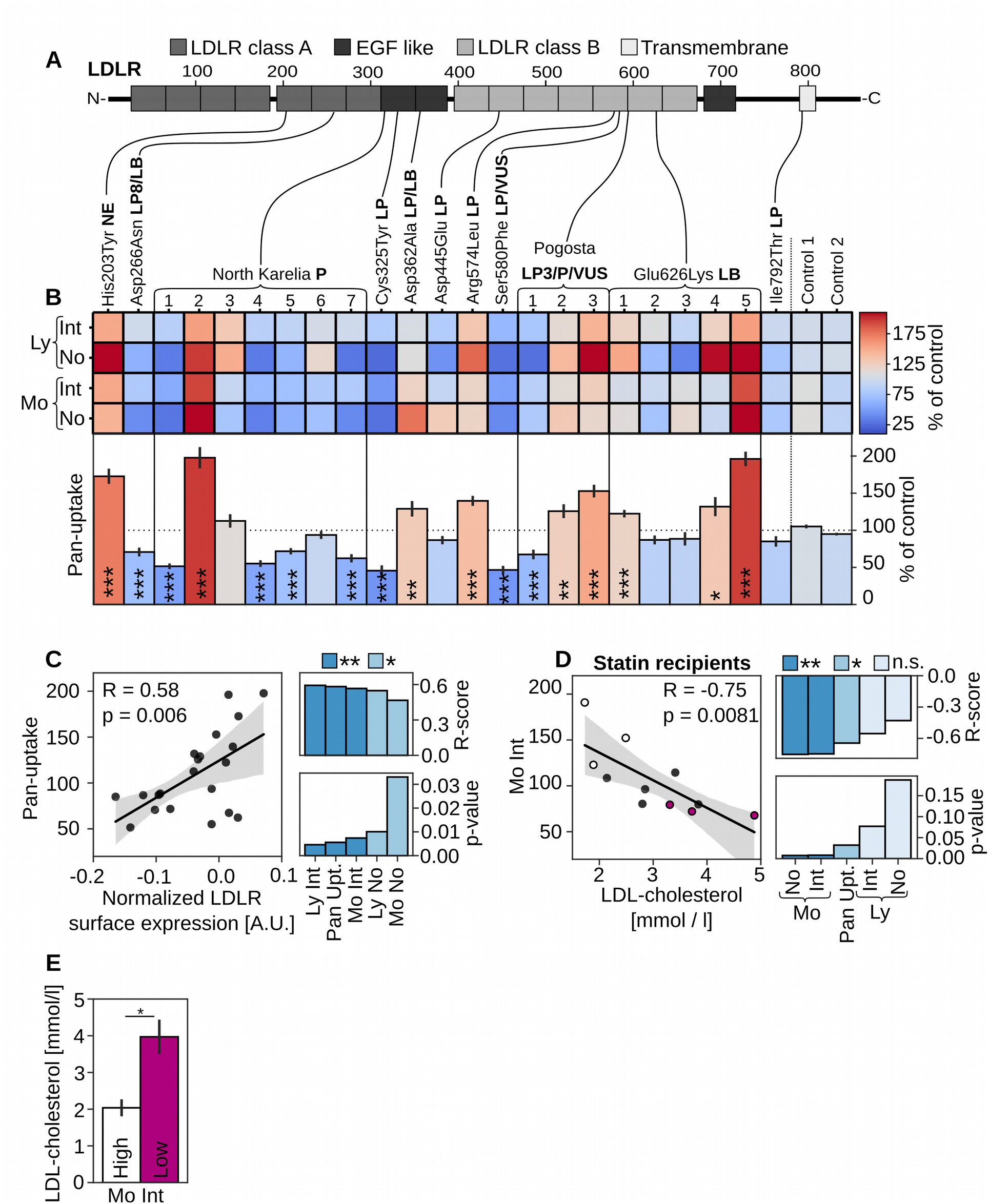
Heterogeneous LDL uptake and LDLR surface expression in He-FH patients’ monocytes. A) Schematic presentation of LDLR mutations included in this study together with their pathogenicity status from ClinVar and LOVD databases. (P = pathogenic, LP = likely pathogenic, LB = likely benign, VUS = variant of unknown significance. B) Quantification of monocyte (Mo) and lymphocyte (Ly) cellular DiI-LDL intensities (Int), organelle numbers (No) and pan-uptake normalized to two controls (100%); two to three independent experiments, each with duplicate or quadruplicate wells per patient (8-16 wells per patient for pan-uptake), Cys325Tyr and Ser580Phe were described in (Figure 1G, H). Significant changes to control two were calculated with Welch’s t-test. c) Correlation of pan-uptake and monocyte LDLR surface expression, including R- and p-values for all uptake scores; n = 21 patients. D) Correlation of monocyte DiI-LDL intensities (Mo Int) with circulating LDL-c for heterozygous FH patients on statin monotherapy, including R- and p-values for all uptake scores. E) LDL-c concentration for 3 patients with the highest (high) and lowest (low) monocyte mean DiI-LDL intensity (Mo Int) as in D. Grey areas in scatter plots indicate 95% CI, *p<0.05, ** p<0.01, *** p<0.001.

Interestingly, the pan-uptake score showed a tendency for lower values in FH-North Karelia carriers as compared to those carrying the likely pathogenic FH-Pogosta and likely benign Glu626Lys variants (**Suppl. Figure 2B**). This is in agreement with higher LDL-c concentrations in FH-North Karelia patients^19^. While LDL uptake did not correlate with circulating LDL-c for the entire study group (**Suppl. Figure 2C**), this correlation was highly significant for monocyte DiI-Int, DiI-No and the pan-uptake scores for the 11 He-FH patients on statin monotherapy (Mo DiI-Int: R=-0.75, p=0.0081, **Figure 2D**). Notably, three of the individuals with the lowest monocyte DiI-Int had a two-fold higher LDL-c concentration than the three individuals with the highest monocyte DiI-Int (**Figure 2E**), suggesting that the LDL-c lowering effect of statin is reflected by monocyte LDL uptake. This is likely due to the higher LDL uptake capacity of monocytes as compared to lymphocytes (**Figure 1E, F**).

### LDL uptake in non-FH individuals with normal or elevated circulating LDL-c

As most hypercholesterolemia patients do not carry *LDLR* mutations, we also investigated cellular LDL uptake in PBMCs from 20 biobank donors with elevated LDL-c levels (LDL-c >5 mM) (hLDL-c) and from 19 donors with normal LDL levels (LDL-c 2-2.5 mM) (nLDL-c) from the FINRISK population cohort^20^ (**Suppl. Table 2**). DNA sequencing confirmed that common Finnish *LDLR* variants were not present among these subjects.

We quantified DiI-Int and DiI-No for monocyte and lymphocyte populations as well as the pan-uptake score for nLDL-c and hLDL-c individuals. This revealed a large interindividual variation in LDL uptake (**Figure 3A**). Both groups included persons with severely reduced LDL internalization, although the lowest pan-LDL uptake scores were among the hLDL-c individuals (**Figure 3A**). Overall, pan-uptake and Ly DiI-No were reduced in hLDL-c compared to nLDL-c subjects, but the differences were not significant (**Suppl. Figure 3A, B**). Of note, reduced pan-uptake, Mo DiI-Int and Ly DiI-No correlated with increased serum LDL-c levels in the hLDL-c subgroup, but the correlations relied on a single individual with a very high serum LDL-c concentration (pan-uptake: R=−0.49, p=0.028; **Suppl. Figure 3C**).

**Figure 3).**
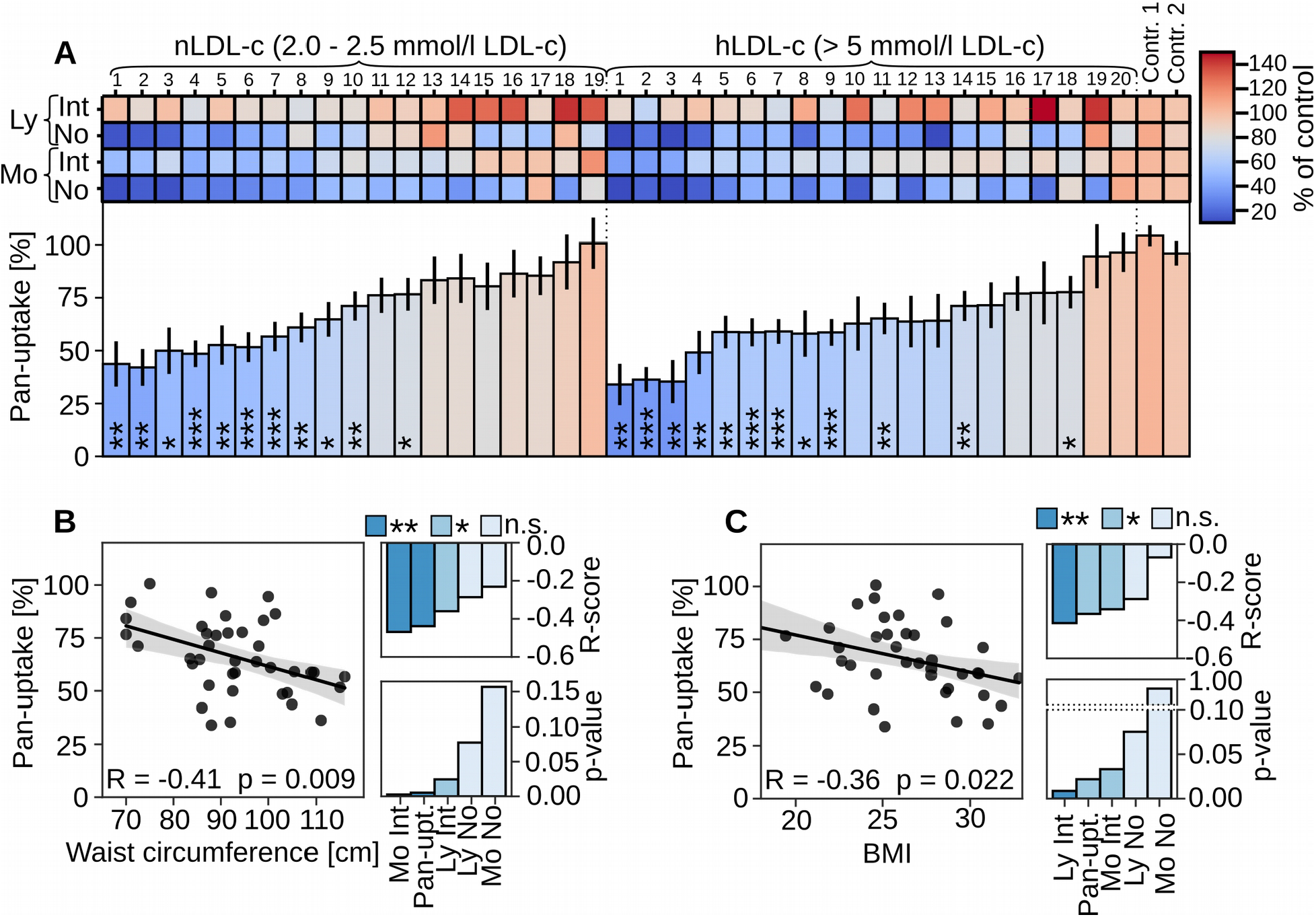
LDL uptake profiles in non-FH individuals with normal and elevated LDL-c. **A**) Quantification of monocyte (Mo) and lymphocyte (Ly) mean DiI-LDL intensities (Int), organelle numbers (No) and pan-uptake after lipid starvation, normalized to control standards; duplicate wells per patient (eight wells per patient for pan-uptake). Significant changes to control two were calculated with Welch’s t-test. **B**) Correlation of pan-uptake with waist circumference and **C**) with body mass index (BMI), including R- and p-values for all uptake scores. n = 39. Grey areas in scatter plots indicate 95% CI. *p<0.05, ** p<0.01, *** p<0.001.

To investigate additional factors influencing the interindividual variability in cellular LDL uptake, we analyzed correlations to two obesity indicators, body mass index (BMI) and waist circumference. Strikingly, reduced pan-uptake, as well as Mo DiI-Int and Ly DiI-Int correlated with increased waist circumference (pan-uptake: R=−0.42, p=0.009; **Figure 3B**). Lower pan-uptake, Ly DiI-Int and Mo DiI-Int also correlated with elevated BMI (pan-uptake: R=−0.36, p=0.022; **Figure 3C**).

### Assessment of cellular lipid storage and mobilization in leukocytes

Cells store excess lipids in LDs and this is related to lipid uptake: When peripheral cells have sufficient lipids available, they typically exhibit LDs and in parallel, lipid uptake is downregulated. We therefore also included the staining of LDs in the automated analysis pipeline (**Figure 1A**). Staining of PBMCs in lipid rich conditions (CM) with the well-established LD dye LD540^21^ revealed that lymphocytes and monocytes displayed LDs in a heterogenous fashion (**Figure 4A**), with lymphocytes showing fewer LD positive cells and fewer LDs per cell than monocytes (**Figure 4B, C**). We then visualized the changes in LD abundance upon overnight lipid starvation in lipoprotein deficient medium (LP) (**Figure 4B-F**). This resulted in a pronounced decrease in lipid deposition: In CM, 9% of lymphocytes and 25% of monocytes contained LDs, but upon lipid starvation, these were reduced to 6% (Ly) and 12% (Mo) (**Figure 4D**).

**Figure 4).**
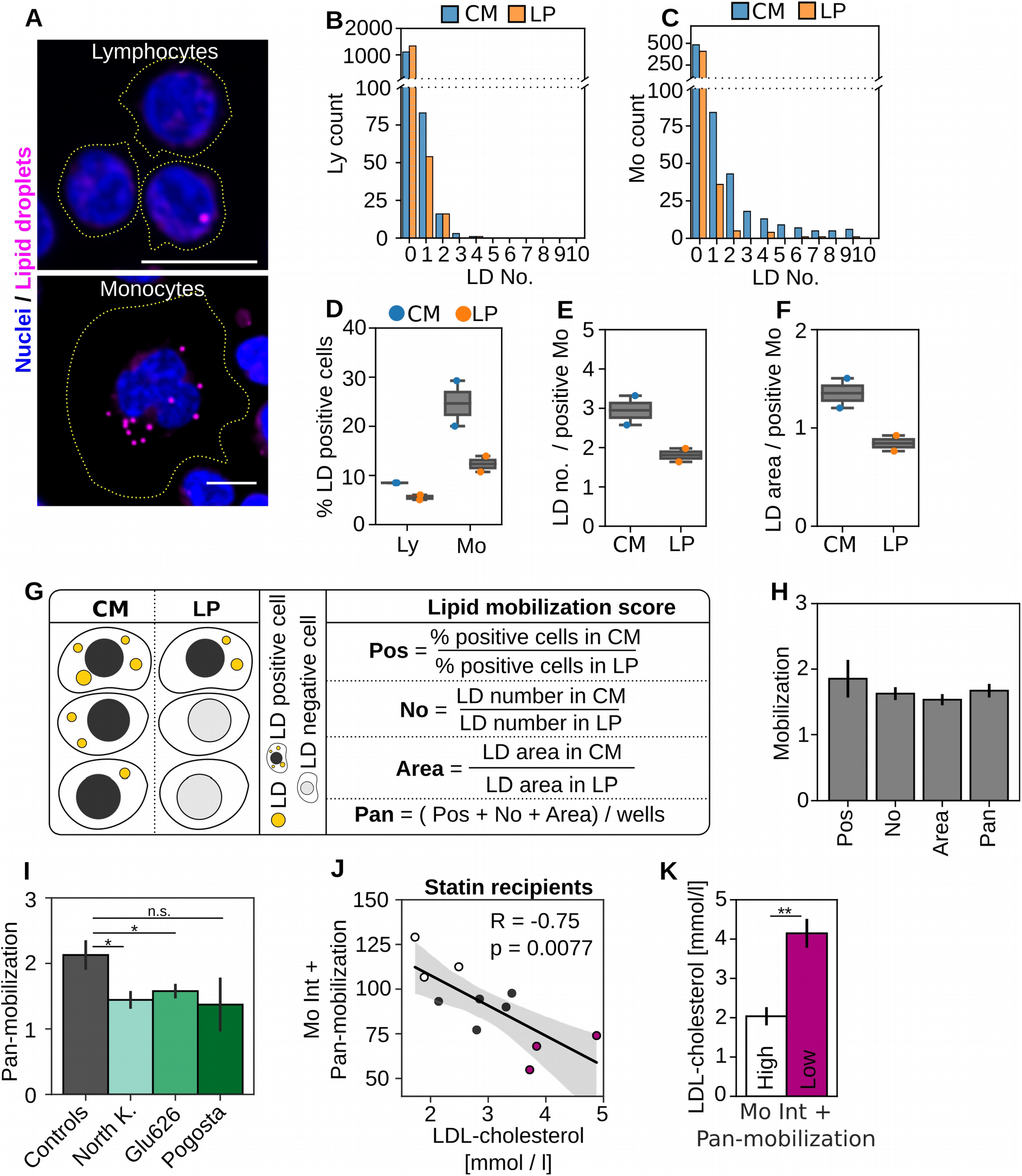
Lipid mobilization assay. **A**) Representative images showing lipid droplets (LDs) in lymphocyte and monocyte populations after treatment with control medium, scale bar = 10μm. **B**) Histogram for cellular LD counts in lymphocyte and (**C**) monocyte populations after treatment with control medium (CM) and lipid starvation (LP) from a single well. **D**) Quantification of LD positive cells in lymphocytes (Ly) and monocytes (Mo) upon treatment with control medium (CM) and lipid starvation (LP); representative of three independent experiments, each with duplicate wells per patient and treatment. **E**) LD counts and (**F**) total LD area in LD positive monocytes quantified for the same experiment as in (**D**). **G**) Schematic presentation of the lipid mobilization score. Upon lipid starvation, the fraction of LD positive monocytes (LD-Pos), their total LD area (LD-Area) and LD numbers (LD-No) are decreasing. Mobilization scores are calculated by dividing the amount of LD-Pos, LD-No or LD-Area in CM with the respective quantifications after lipid starvation. Pan-mobilization is the average of LD-Pos, LD-No and LD-Area mobilization scores from individual wells. **H**) Lipid mobilization scores for one control; n = 6 wells from three independent experiments, (18 wells for pan-mobilization), ± SEM. **I**) Pan-mobilization for controls (combined control one and two from five experiments), FH-North-Karelia (n = 7), FH-Pogosta (n = 3) and FH-Glu626 (n = 5). **J**) Correlation of combined monocyte mean DiI-LDL intensities (Mo Int) and pan-mobilization with circulating LDL-c. **K**) LDL-c concentration for 3 patients with the highest (high) and lowest (low) combined score as in **J**. *p < 0.05, **p < 0.01.

Due to the lower LD abundance in lymphocytes, we focused on monocytes and defined three readouts for them: 1) Percentage of LD-positive cells (LD-Pos), 2) Cellular LD number in LD-Pos (LD-No) and 3) Total cellular LD Area in LD-Pos (LD-Area). On average, LD-Pos cells showed 2.9 LDs in lipid rich conditions and 1.8 LDs upon lipid starvation (**Figure 4E**), while the total LD area decreased from 1.35 μm^2^ in lipid rich conditions to 0.8 μm^2^ upon lipid starvation (**Figure 4F**).

When quantifying LD parameters from several subjects, we observed substantial differences between individuals in how LDs changed upon starvation. To systematically quantify these differences, we established a parameter, lipid mobilization score, that reflects how efficiently cellular lipid stores are depleted under lipid starvation (**Figure 4G**). Lipid mobilization scores were calculated for each of the LD readouts, LD-Pos, LD-No and LD-Area, by dividing the results obtained in lipid rich conditions with those obtained after lipid starvation (**Figure 4G**). Furthermore, we established a pan-mobilization score by averaging LD-Pos, LD-No and LD-Area scores (**Figure 4G, H**), with LD-Pos providing the highest mobilization score but also the highest variability (**Figure 4H**).

To further assess the reliability of the LD mobilization parameters, we determined their intraindividual variation using the same samples as for analyzing intraindividual variation of DiI-LDL uptake (**Suppl. Figure 1I, J**). This showed a modest intraindividual variation for the lipid mobilization scores (**Suppl. Figure 4A**), which an average of 8% for pan-mobilization, 10% for LD-Pos, 11% for LD-No and 13% for LD-Area (**Suppl. Figure 4B**).

### Cellular lipid mobilization in He-FH patients

When lipid mobilization was analyzed from the He-FH samples of the METSIM cohort, we found that the pan-mobilization score was significantly reduced in He-FH individuals carrying the FH-North Karelia and Glu626Lys variants (**Figure 4I**). This suggests that defective LDLR function may be accompanied by reduced lipid mobilization. We also studied whether the combination of a lipid mobilization score with LDL uptake improves identification of statin recipients with high residual LDL-c concentrations. Several of the patients with intermediate and high LDL-c showed low monocyte DiI-LDL intensities in a narrow range (**Figure 2D**). When monocyte DiI-Int was combined with the pan-mobilization score, larger differences between patients were observed, providing a better separation of individuals with high and intermediate LDL-c (**Figure 4J**). Moreover, the difference in LDL-c concentration between the three individuals with the highest vs. lowest score was more significant than when using monocyte DiI-Int alone (**Figure 4K vs. Figure 2E**). This suggests that the combined LDL uptake and lipid mobilization assays may help to better pinpoint those He-FH cases that remain refractory to statin monotherapy.

### Cellular lipid mobilization is reduced in non-FH hypercholesterolemia patients and correlates with LDL uptake

We then investigated whether monocytes from nLDL-c and hLDL-c biobank donors displayed differences in lipid mobilization. Analogously to LDL uptake, we observed a large variability for the pan- and individual mobilization scores in this cohort (**Figure 5A**). Interestingly, pan-mobilization, LD-No and LD-Area were significantly reduced in the hLDL-c compared to nLDL-c subjects (**Figure 5A, B, Suppl. Figure 5A, B**). This prompted us to scrutinize whether lipid mobilization correlates with LDL uptake related parameters or obesity indicators in this cohort. All mobilization scores correlated positively with the pan-uptake score (R=0.42, p=0.0095 for pan-mobilization; **Figure 5C**). Furthermore, pan-, LD-No and LD-Area mobilization scores correlated negatively with total cholesterol, apo-B concentrations (**Suppl. Figure 5C, D**) and with age (R=−0.38, p=0.019 for pan-mobilization; **Figure 5D)**.

**Figure 5).**
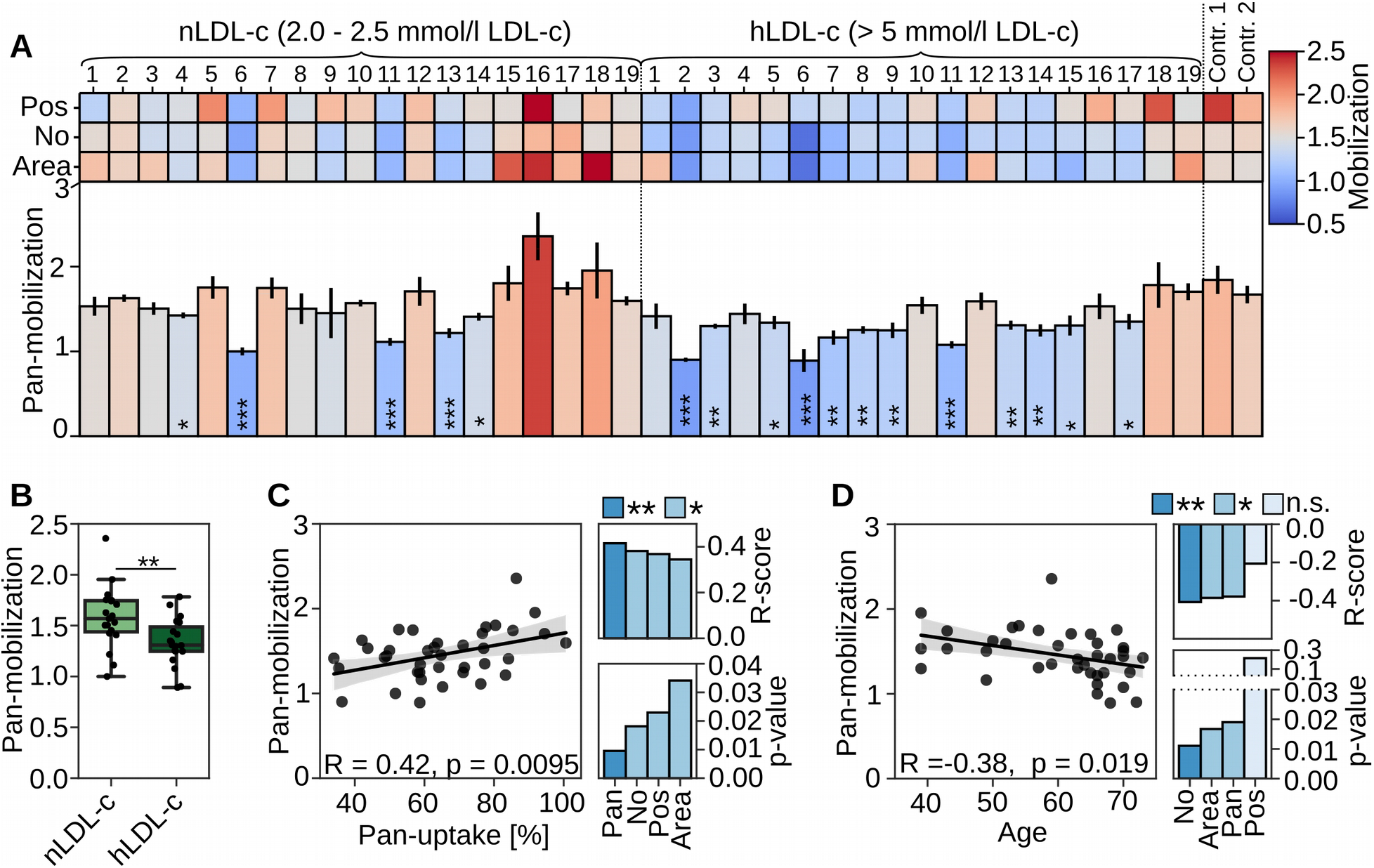
Monocyte lipid mobilization correlates with LDL uptake and is reduced in subjects with elevated LDL-c. A) Mobilization scores (Pos, LD-No, LD-Area and pan-mobilization) in monocytes from controls (nLDL-c, LDL-c 2-2.5 mmol/l) and individuals with elevated LDL-c (hLDL-c, LDL > 5 mmol/l) sorted according to the pan-uptake score (Figure 3A); duplicate wells per patient (six wells per patient for pan-mobilization). Significant changes to control two were quantified with Welch’s t-test. B) Box plot of pan-mobilization for nLDL-c and hLDL-c subgroups; nLDL-c n = 19, hLDL-c n = 19. ** p < 0.01, Students t-test. Correlation of pan-mobilization with pan-uptake (C) and age (D), including R- and p-values for all mobilization scores. Grey areas in scatter plots = 95% CI. * p<0.05, * p<0.01, *p<0.001.

### Hybrid scores of genetic and functional cell-based data show improved association with hypercholesterolemia

The hLDL-c biobank donors of the FINRISK population cohort displayed an increased LDL-c polygenic risk score (LDL-PRS) (**Figure 6A**). LDL-PRS did not correlate with LDL uptake or lipid mobilization (**Suppl. Figure 6A, B**), suggesting that LDL-PRS and cellular LDL uptake monitor in part distinct processes. Interestingly, combination of LDL-PRS with pan-uptake reduced the variation and made it easier to discriminate the nLDL-c and hLDL-c groups, providing an eight times better p-value as compared to LDL-PRS only (**Figure 6B**). Furthermore, combination of the pan-mobilization score with LDL-PRS drastically improved the discrimination between groups (**Figure 6C**) and combining all three parameters, i.e. LDL-PRS, pan-uptake and pan-mobilization, provided the best discrimination power and lowest p-value (**Figure 6D**). To estimate the association of LDL-PRS and novel hybrid scores with elevated LDL-c (>5 mmol/l), we calculated the odds ratio (OR) for elevated LDL-c by comparing individuals with the highest 30% of the score to the remaining subjects. Combining LDL-PRS with either pan-uptake or pan-mobilization doubled the OR and using a hybrid score combining all three readouts resulted in a five-fold higher OR (**Figure 6E**).

**Figure 6).**
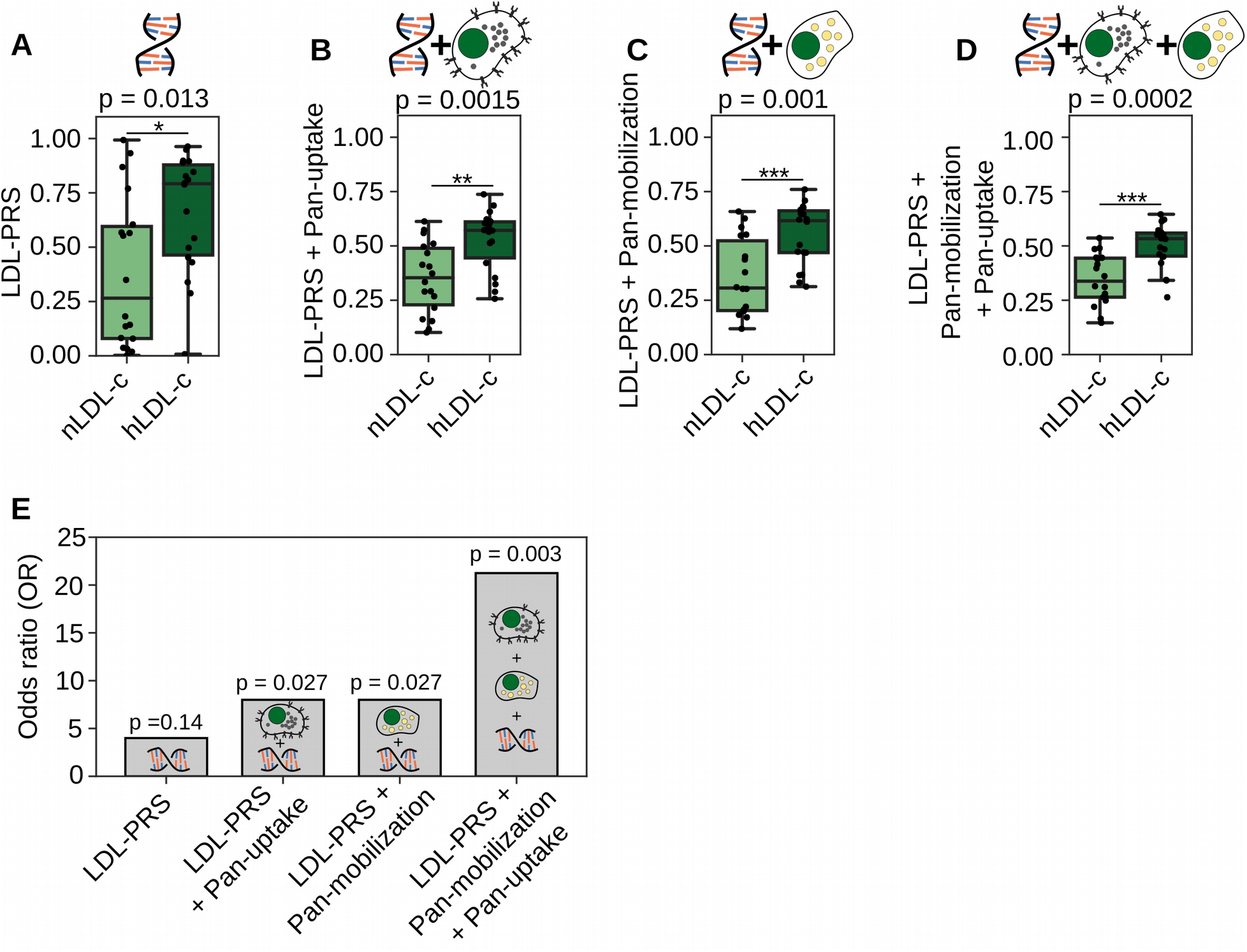
Hybrid scores combining genetic and functional cell-based data show improved association with hypercholesterolemia. A) Box plot of a polygenic risk score for high LDL-c levels (LDL-PRS) for nLDL-c (2-2.5 mmol/l LDL-c) and hLDL-c (>5 mmol/l LDL-c) subgroups. B) Box plot for double hybrid scores combining LDL-PRS and pan-uptake or pan-mobilization (C) into a single score. D) Box plot for a triple hybrid score consisting of LDL-PRS, pan-uptake and mobilization. nLDL-c n = 18, hLDL-c n = 19, * p<0.05, ** p<0.01, *** p<0.001; Welch’s t-test. E) Odds ratio (OR) for 30% of the individuals with the highest LDL-PRS, double or triple hybrid scores and the remaining subjects, calculated with the Fisher’s exact probability test.

## Discussion

In this study, we established a multiplexed analysis pipeline to quantify lipid uptake and mobilization in primary leukocytes and used it to analyze over 300 conditions (combinations of assays and treatments) from 65 individuals. The automated cell handling, staining and imaging procedures enable high-throughput applications. Key advantages of the method are: 1) Large-scale internal standards allow comparison of experimental results over time, 2) Automated cell quantification avoids researcher bias, increasing reliability of results, 3) Semi-automated workflow can be scaled to increase throughput, 4) Cell immobilization on coated surfaces allows flexibility in sample handling and facilitates automation, 5) Lymphocyte- and monocyte-enriched cell populations can be detected based on cell spreading on coated surfaces, 6) Subcellular resolution enables quantification of internalized LDL and LDs, yielding new scores derived from them. While the first two aspects are or can be readily included in conventional flow cytometry based LDL uptake assays, the latter four rely on a high-content high-resolution imaging platform.

Several of the observations made using this analysis pipeline are supported by previous findings obtained using manual assays, thereby validating our results. We showed that monocytes display higher LDL uptake activities than lymphocytes, in accordance with previous findings^16^. The highly variable LDL uptake observed by us between individuals, including He-FH patients with identical *LDLR* mutations, also agrees with earlier reports^10–12^. Furthermore, we observed an association of low cellular LDL uptake with increased circulating LDL-c in He-FH patients on statin monotherapy, in line with studies utilizing radiolabelled LDL^22–25^. However, this finding was not readily reproduced by using fluorescently labelled LDL particles in lymphocytes^26,27^. Indeed, our results indicate that monocytes provide an improved detection window and a better correlation between cellular LDL uptake and circulating LDL-c.

We also found that reduced LDL uptake correlated with increased BMI and waist circumference, two obesity indicators. Metabolic syndrome is typically linked to dyslipidemia characterized by decreased high-density lipoprotein cholesterol (HDL-c), elevated LDL-c with increased small dense LDL particles and increased plasma triglycerides^37^. Our results suggest, that besides VLDL overproduction and defective lipolysis of TG-rich lipoproteins^1^, reduced LDL clearance may contribute to dyslipidemia in overweight individuals. This fits with the observed reduction of LDLR expression in obese subjects^38^.

Moreover, we employed the platform to quantify cellular LDs, established a new parameter termed lipid mobilization score, and demonstrated its ability to provide additional data on individual differences on lipid handling. Lipid mobilization correlated with LDL uptake, implying that efficient removal of stored lipids was typically paralleled by efficient lipid uptake. Moreover, combining monocyte LDL uptake and lipid mobilization data facilitated the detection of He-FH cases that remained hypercholesterolemic on statin. In the FINRISK population cohort, lipid mobilization outperformed LDL uptake in distinguishing individuals with high (>5 mmol/l) and normal LDL-c (2-2.5 mmol/l), with impaired lipid mobilization associating with elevated LDL-c. Hence, lipid mobilization shows potential to highlight additional aspects of cellular lipid metabolism underlying hypercholesterolemia in individual patients.

Polygenic risk scores (PRS) provide tools for cardiovascular risk profiling and are increasingly included in clinical care guidelines of hypercholesterolemia^1,28^. We found that the hypercholesterolemia subjects of the FINRISK cohort had an increased LDL-PRS, but this did not correlate with LDL uptake or lipid mobilization, arguing that the cell-based parameters cover in part different territories than PRS. In agreement, the combination of LDL-uptake, lipid mobilization and LDL-PRS improved the segregation of hyper- and normocholesterolemic subjects. An increased LDL-PRS is associated with a higher incidence of coronary artery disease^5^. We therefore anticipate that the cell-based assays may provide additional information for future integrated CVD risk calculations. These, in turn, might facilitate the detection of hypercholesterolemia risk at younger age when clinical manifestations are not yet overt, enabling faster initiation of treatment and improved disease prevention^29^.

In summary, the automated analysis platform established here enables systematic assessment of cellular lipid trafficking in accessible primary cell samples of human origin. Besides hypercholesterolemia, this approach can be useful in other metabolic disorders, as well as diseases not previously linked to cellular lipid imbalance. As an example of the latter, we recently uncovered aberrant LD size distribution in MYH9-related disease patient neutrophils using quantitative imaging^30^.

## Limitations of the study

We analyzed 65 individuals as a proof-of-concept for the analysis platform. Whilst this outperforms most previous studies measuring lipid uptake in primary cells, further validation in larger study groups will be required. Such studies will be feasible due to the high automation level of the platform, enabling processing of samples from several thousand subjects per year.

Regarding the cellular origin of hypercholesterolemia, we infer parameters related to whole body metabolism and in particular liver function from PBMCs. Evidently, primary hepatocytes would provide more direct information, but are not accessible on a routine basis. PBMCs are easily obtained from standard blood collections. Moreover, our data demonstrate that PBMC derived parameters can correlate with readouts deriving from the whole body level.

Currently, the analysis platform is set up to quantify two cellular parameters, LDL uptake and lipid storage in droplets. In the future, the utility of the platform can be further extended by the inclusion of additional fluorescence based readouts amenable to high-content imaging and quantification.

## Acknowledgements

We thank Anna Uro for technical assistance, HiLIFE and Biocenter Finland supported Helsinki BioImaging infrastructures for help with microscopy, Katariina Öörni for help with LDL preparation and Abel Szkalisity for help with image analysis. We thank THL Biobank for providing samples and data for this study (study no: 2016_15, 2016_117 and 2018_15) and all biobank donors for their participation in biobank research.

This study was supported by The Academy of Finland (grants 282192, 284667, 307415 to EI; 321428 to ML; 310552 to LP; 328861, 325040 to SP; and 312062, 316820 to SR), Sigrid Juselius Foundation (grants to EI, ML SR), University of Helsinki (grant to KK; Faculty of Medicine early-career investigator grant to SP; HiLIFE Fellow grant to EI), Finnish Foundation for Cardiovascular Research and University of Helsinki HiLIFE Fellow and Grand Challenge grants, and H2020-INTERVENE (101016775) to SR, Fondation Leducq (grant 19CVD04 to EI), MIUR of Italy (project cod. PON03PE_00060_7, grant to CEINGE, G.F.), LENDULET-BIOMAG Grant (2018-342), H2020-discovAIR (874656), and Chan Zuckerberg Initiative (seed networks for the HCA-DVP) to PH. Ida Montin Foundation (grant to PR), Doctoral Programme in Population Health, University of Helsinki (grant to PR); Emil Aaltonen Foundation (grant to PR).

## Author Contributions

S.G.P and E.I. designed the study and developed the concept. S.G.P, I.B., K.K., I.H., and P.R. performed experiments. S.G.P, I.B., K.K. I.H., P.R., M.M.I., S.R. and E.I. analyzed data and interpreted results, A.K., M.D.T., A.S.F., G.F., J.K. and M.L. provided patient samples and clinical data. L.P. and P.H. established image analysis and processing tools. S.G.P and E.I. wrote the manuscript. All authors reviewed and revised the manuscript.

## STAR Methods

### Materials

Lipoprotein deficient serum (LPDS) was obtained from fetal bovine serum by ultracentrifugation as described^31^. For DiI-LDL, we first prepared fresh LDL from human plasma samples (Finnish Red Cross permit 39/2016) by density centrifugation^32^ and then labelled LDL with 1,1’-dioctadecyl-3,3,3’,3’-tetramethyl-indocarbocyanine perchlorate (DiI) as described^33^. 4′,6-diamidino-2-phenylindole (DAPI), Poly-D lysine (PDL) and Histopacque Premium were obtained from Sigma. DiI, anti-mouse Alexa 568, HCS CellMask Deep Red and HCS CellMask Green were obtained from Thermo Fisher. Mouse anti-LDLR (clone 472413) was from R&D systems.

### Peripheral blood mononuclear cells (PBMC) and blood samples

All blood samples were collected in accordance with the declaration of Helsinki regarding experiments involving humans. He-FH patients were identified in the Metabolic Syndrome in Men study (METSIM)^18^ and blood samples obtained during patient follow-up. Two He-FH patients (Cys325Tyr and Ser580Phe) for which we obtained PBMC and EBV lymphoblast samples were described previously^34^. PBMC samples from the Finnish population survey, FINRISK 2012, and the donor linked data (including genotypes) were obtained from THL Biobank (www.thl.fi/biobank) and used under the Biobank agreements no 2016_15, 2016_117 and 2018_15. The FINRISK 2012 study groups consisting of donors with elevated LDL-c levels (LDL > 5 mM, hLDL-c) and normal levels (LDL-c 2.0-2.5 mM, nLDL-c) were age, gender and BMI matched. The donors in neither of groups had cholesterol lowering medication by the time of sampling, and based on a food frequency questionnaire, did not receive an elevated proportion of energy intake as saturated or trans-fat. Buffy coat samples from healthy blood donors were obtained from the Finnish Red Cross (permit 392016). Three healthy volunteers donated blood samples on two consecutive days after overnight fasting, to assess the intraindividual variation of LDL uptake and lipid mobilization.

### Cell culture

Control EBV lymphoblasts (GM14664) were obtained from Coriell Cell Repository and cultured in RPMI-1640 supplemented with 15% FBS, penicillin/streptomycin (100 U/ml each) and 2 mM L-Glutamine. For continuous culturing of EBV lymphoblasts, 3×10^6^ cells were transferred to 5 ml of fresh medium once a week. Cells were cryopreserved in 70% PBMC medium (RPMI-1640, penicillin/streptomycin, 2 mM L-glutamine, 1 mM sodium pyruvate, and 1 mM HEPES), 20% FBS and 10% DMSO.

### PBMC isolation

Blood or buffy coat samples were mixed 1:1 with phosphate buffered saline (PBS) including 2.5 mM EDTA (PBS-E). The blood mixture was gently layered over Histopaque Premium (1.0073, for mononuclear cells) and centrifuged 40 min at 400 g. The PBMC cell layer was removed, transferred to a new 15 ml reaction tube and mixed with PBS-E. Cells were centrifuged at 400 g for 10 min and incubated in 2 ml of red blood cell lysis buffer for 1 min (155 mM NH_4_Cl, 12 mM NaHCO_3_, 0.1 mM EDTA). 10 ml of PBS-E was added and cells were pelleted and washed with PBS-E. Then cells were resuspended in 5 ml PBMC medium (RPMI-1640, penicillin/streptomycin, 2 mM L-glutamine, 1 mM sodium pyruvate, and 1 mM HEPES), counted, pelleted and cryopreserved.

### Cell treatments, DiI-LDL uptake, transfer to imaging plates and fixation

Cryopreserved EBV lymphoblasts or PBMCs were thawed in PBMC medium, and centrifuged at 400 g for 10 min. The cells were resuspended in PBMC medium and transferred to a well of a 96 well plate (200000 cells per well), containing FBS (10% final concentration) or LPDS (5% final concentration) and incubated for 24 h. Cells were then incubated with freshly thawed DiI-LDL at 30 μg / ml final concentration for 1 h at 37°C, which yielded an optimal signal intensity at a linear detection range in PBMCs. Subsequently, cells were transferred to conical 96 well plates and centrifuged at 400 g for 10 min. Using a robotic platform (Opentrons, New York, USA) medium was removed and cells were resuspended in PBMC medium. Cells were centrifuged, automatically resuspended in PBMC medium and transferred to PDL coated 384 well high-content imaging plates (approximately 40 000 cells/well, a density where individual cells are not on top but close to each other). The robotic resuspension ensured homogenous cell adhesion to the imaging plates. After 30 min of incubation at 37°C cells were automatically fixed with 4% paraformaldehyde in 250 mM HEPES, 1 mM CaCl_2_, 100 μM MgClM MgCl_2_, pH 7.4 and washed with PBS. For lipid droplet and LDLR surface stainings, cells were directly transferred to PDL coated 384 well high-content plates, adhered, automatically fixed and washed with PBS.

### Lipid droplet analyses

Cells were processed as described before^27^ with the following changes: Fixed cell samples were automatically stained with 1 μg/ml LD540 (Princeton BioMolecular Research) and 5 μg/ml DAPI. 3D stacks of optical slices were acquired automatically either with a Nikon Eclipse Ti-E inverted microscope equipped with a 40 × Planfluor objective with NA 0.75 and 1.5 zoom; duplicate wells, each with six image fields per patient, or with a PerkinElmer Opera Phenix High Content Imaging system with a 63x water immersion objective, NA 1.15; duplicate wells, each with 14, 16 (two wells combined) or 24 (two wells combined) image fields. Image stacks were automatically deconvolved either with Huygens software (Scientific Volume Imaging, b.v.) or a custom-made Python tool based on the open-source tools PSF generator^35^ and deconvolution lab^36^. Maximum intensity projections were made from the deconvolved image stacks with custom Python tools. Automated quantification of lipid droplets was performed as described previously^30,37,38^.

### LDLR surface staining

All staining procedures were performed automatically. Fixed cells were quenched with 50 mM NH4Cl for 15 min and washed twice with PBS. Cells were incubated with block solution (PBS, 1% BSA) for 10 min followed by staining with mouse anti-LDLR in block solution for 60 min. Cells were washed three times with PBS followed by incubation with secondary antibody solution (anti-mouse-Alexa 568, DAPI 5 μg/ml and HCS CellMask Green stain 0.25 μg/ml) for 45 min at room temperature. Cells were washed with PBS and 3D stacks of optical slices were acquired for DAPI (nuclei), CellMask Green (cytoplasm), Alexa 568 (LDLR surface) and Alexa 640 (background) channels using an Opera Phenix high-content imaging system with a 40x water immersion objective NA 1.1; quadruplicate wells, each with seven image fields per patient. LDLR surface and background images were automatically deconvolved with our custom build Python deconvolution tools and maximum intensity projections were made. The resulting images were automatically analysed with CellProfiler^39^. LDLR surface intensities were background subtracted for each individual cell and normalized by subtracting mean LDLR surface intensities from the two controls, which were included in each imaging plate.

### Quantification of DiI-LDL uptake

DiI-LDL labeled, and fixed cells (see section cell treatments) were automatically processed with a robotic platform (Opentrons). Cells were stained with 5 μg/ ml DAPI and 0.5 μg/ml HCS CellMask Deep Red and image stacks for three channels, DAPI (nuclei), DiI-LDL and CellMask Deep Red (cytoplasm) were acquired. Automated microscopy and single cell quantifications with CellProfiler were performed as described in the section LDLR surface staining; Quadruplicate wells, each with 7 image fields for heterozygous FH patients; duplicate wells, each with 13 image fields for FINRISK subjects. Plate effects were determined with control samples and corrected for in the individual experiments.

### LDL-c polygenic risk score (LDL-PRS)

Genotyping of FINRISK2012 samples has been previously described^5^ We calculated the LDL PRS using the LDpred method based on both an European genome-wide association study (GWAS) meta-analysis with 56945 samples and the previously published PRS by *Talmud* et al.^4,40^. The PRS calculation is described in detail in the Supplemental materials. LDL uptake and lipid mobilization parameters were normalized to a range from 0 to 1 to generate uptake and mobilization scores. Hybrid scores represent the average of LDL-PRS and uptake and/or mobilization scores which were normalized to a range from 0 to 1.

### Data analysis

Segmented images from CellProfiler underwent routine visual controls to verify cell identification and filter out potential imaging artifacts. Then, lymphocytes and monocytes were detected based on the size of the cytoplasm (Ly <115 μm^2^, Mo >115 μm^2^) (See Suppl. Figure 1). We averaged the cellular mean DiI-LDL intensities and organelle counts for each cell population and well and normalized them to the average of both controls included in each plate, set to 100%. For LD quantifications we first selected monocytes with at least one LD. We then averaged cellular LD number and total LD area (LD number x LD size) for each well. For lipid mobilization we first averaged the control medium results for LD-Pos, LD-No, and LD-area from duplicate wells and then divided these by the respective per well results after lipid starvation. We used Python (Python Software Foundation, www.python.org) with the following packages to perform the single cell data analysis (Pandas, Numpy, Scipy, Matplotlib^41^, Seaborn^42^). For statistical significance testing we utilized aggregated single cell data at the level of individual wells (n = number of wells per treatment and patient). First, we performed Levene’s test to assess the equality of sample variation. For equal sample amounts and variance, we carried out a two-tailed Student’s t-test. For unequal samples or variance, we utilized Welch’s t-test. For correlations we first performed a linear regression of the two measurements and then calculated a two-sided p-value for a hypothesis test whose null hypothesis is that the slope is zero, using Wald Test with t-distribution of the test statistic. Fisher’s exact probability test was used to calculate the odds ratio. Among the FINRISK2012 hLDL-c subgroup there is one individual with a serum LDL-c of 10.1 mmol / l. We performed a sensitivity analysis by removing this subject from our analysis, to verify that the major conclusions of this study are not affected by this individual.

### Data and code availability

The data supporting the findings of this study are available from the authors upon request. Genetic data for the subjects of the FINRISK cohort study is available from the THL Biobank (https://thl.fi/en/web/thl-biobank). Custom python tools for image processing and deconvolution can be accessed via: https://github.com/lopaavol/Oputils. Software tools for lipid droplet detection have been described previously^38^ and are available via: https://bitbucket.org/szkabel/lipidanalyzer/get/master.zip

## References

1. Borén, J. et al. Low-density lipoproteins cause atherosclerotic cardiovascular disease: pathophysiological, genetic, and therapeutic insights: a consensus statement from the European Atherosclerosis Society Consensus Panel. Eur. Heart J. (2020) doi:10.1093/eurheartj/ehz962.

2. Abul-Husn, N. S. et al. Genetic identification of familial hypercholesterolemia within a single U.S. health care system. Science 354, aaf7000 (2016).

3. Khera, A. V. et al. Diagnostic Yield and Clinical Utility of Sequencing Familial Hypercholesterolemia Genes in Patients With Severe Hypercholesterolemia. Journal of the American College of Cardiology 67, 2578–2589 (2016).

4. Talmud, P. J. et al. Use of low-density lipoprotein cholesterol gene score to distinguish patients with polygenic and monogenic familial hypercholesterolaemia: a case-control study. The Lancet 381, 1293–1301 (2013).

5. Ripatti Pietari et al. Polygenic Hyperlipidemias and Coronary Artery Disease Risk. Circulation: Genomic and Precision Medicine 13, e002725 (2020).

6. Ray, K. K. et al. EU-Wide Cross-Sectional Observational Study of Lipid-Modifying Therapy Use in Secondary and Primary Care: the DA VINCI study. European Journal of Preventive Cardiology (2020) doi:10.1093/eurjpc/zwaa047.

7. Snijder, B. et al. Image-based ex-vivo drug screening for patients with aggressive haematological malignancies: interim results from a single-arm, open-label, pilot study. Lancet Haematol 4, e595–e606 (2017).

8. Romano, M. et al. Identification and functional characterization of LDLR mutations in familial hypercholesterolemia patients from Southern Italy. Atherosclerosis 210, 493–496 (2010).

9. Benito-Vicente, A. et al. Validation of LDLr Activity as a Tool to Improve Genetic Diagnosis of Familial Hypercholesterolemia: A Retrospective on Functional Characterization of LDLr Variants. International Journal of Molecular Sciences 19, 1676 (2018).

10. Urdal, P., Leren, T. P., Tonstad, S., Lund, P. K. & Ose, L. Flow cytometric measurement of low density lipoprotein receptor activity validated by DNA analysis in diagnosing heterozygous familial hypercholesterolemia. Cytometry 30, 264–268 (1997).

11. Tada, H. et al. A novel method for determining functional LDL receptor activity in familial hypercholesterolemia: Application of the CD3/CD28 assay in lymphocytes. Clinica Chimica Acta 400, 42–47 (2009).

12. Thedrez Aurélie et al. Homozygous Familial Hypercholesterolemia Patients With Identical Mutations Variably Express the LDLR (Low-Density Lipoprotein Receptor). Arteriosclerosis, Thrombosis, and Vascular Biology 38, 592–598 (2018).

13. Ikonen, E. Cellular cholesterol trafficking and compartmentalization. Nat. Rev. Mol. Cell Biol. 9, 125–138 (2008).

14. Luo, J., Yang, H. & Song, B.-L. Mechanisms and regulation of cholesterol homeostasis. Nat Rev Mol Cell Biol 21, 225–245 (2020).

15. Chan, P., Jones, C., Lafrenière, R. & Parsons, H. G. Surface expression of low density lipoprotein receptor in EBV-transformed lymphocytes: characterization and use for studying familial hypercholesterolemia. Atherosclerosis 131, 149–160 (1997).

16. Schmitz G, Brüning T, Kovacs E & Barlage S. Fluorescence flow cytometry of human leukocytes in the detection of LDL receptor defects in the differential diagnosis of hypercholesterolemia. Arteriosclerosis and Thrombosis: A Journal of Vascular Biology 13, 1053–1065 (1993).

17. Piccaluga, P. P., Weber, A., Ambrosio, M. R., Ahmed, Y. & Leoncini, L. Epstein–Barr Virus-Induced Metabolic Rearrangements in Human B-Cell Lymphomas. Front Microbiol 9, (2018).

18. Laakso, M. et al. The Metabolic Syndrome in Men study: a resource for studies of metabolic and cardiovascular diseases. J. Lipid Res. 58, 481–493 (2017).

19. Lahtinen, A. M., Havulinna, A. S., Jula, A., Salomaa, V. & Kontula, K. Prevalence and clinical correlates of familial hypercholesterolemia founder mutations in the general population. Atherosclerosis 238, 64–69 (2015).

20. Borodulin, K. et al. Cohort Profile: The National FINRISK Study. Int J Epidemiol 47, 696–696i (2018).

21. Spandl, J., White, D. J., Peychl, J. & Thiele, C. Live Cell Multicolor Imaging of Lipid Droplets with a New Dye, LD540. Traffic 10, 1579–1584 (2009).

22. Hagemenas F C & Illingworth D R. Cholesterol homeostasis in mononuclear leukocytes from patients with familial hypercholesterolemia treated with lovastatin. Arteriosclerosis: An Official Journal of the American Heart Association, Inc. 9, 355–361 (1989).

23. Hagemenas, F. C., Pappu, A. S. & Illingworth, D. R. The effects of simvastatin on plasma lipoproteins and cholesterol homeostasis in patients with heterozygous familial hypercholesterolaemia. European Journal of Clinical Investigation 20, 150–157 (1990).

24. Gaddi, A. et al. Pravastatin in heterozygous familial hypercholesterolemia: Low-density lipoprotein (LDL) cholesterol-lowering effect and LDL receptor activity on skin fibroblasts. Metabolism 40, 1074–1078 (1991).

25. Sun, X.-M., Patel, D. D., Knight, B. L. & Soutar, A. K. Influence of genotype at the low density lipoprotein (LDL) receptor gene locus on the clinical phenotype and response to lipid-lowering drug therapy in heterozygous familial hypercholesterolaemia. Atherosclerosis 136, 175–185 (1998).

26. Raungaard, B., Brorholt Petersen, J. U., Jensen, H. K. & Færgeman, O. Flow Cytometric Assessment of Effects of Fluvastatin on Low-Density Lipoprotein Receptor Activity in Stimulated T-Lymphocytes from Patients with Heterozygous Familial Hypercholesterolemia. The Journal of Clinical Pharmacology 40, 421–429 (2000).

27. Homma, K. et al. Changes in ultracentrifugally separated plasma lipoprotein subfractions in patients with polygenic hypercholesterolemia, familial combined hyperlipoproteinemia, and familial hypercholesterolemia after treatment with atorvastatin. Journal of Clinical Lipidology 9, 210–216 (2015).

28. Mach, F. et al. 2019 ESC/EAS guidelines for the management of dyslipidaemias: Lipid modification to reduce cardiovascular risk. Atherosclerosis 290, 140–205 (2019).

29. Wiegman, A. et al. Familial hypercholesterolaemia in children and adolescents: gaining decades of life by optimizing detection and treatment. Eur Heart J 36, 2425–2437 (2015).

30. Pfisterer, S. G. et al. Role for formin-like 1-dependent acto-myosin assembly in lipid droplet dynamics and lipid storage. Nature Communications 8, 14858 (2017).

31. Goldstein, J. L., Basu, S. K. & Brown, M. S. Receptor-mediated endocytosis of low-density lipoprotein in cultured cells. Meth. Enzymol. 98, 241–260 (1983).

32. Stephan, Z. F. & Yurachek, E. C. Rapid fluorometric assay of LDL receptor activity by DiI-labeled LDL. J. Lipid Res. 34, 325–330 (1993).

33. Reynolds, G. D. & St Clair, R. W. A comparative microscopic and biochemical study of the uptake of fluorescent and 125I-labeled lipoproteins by skin fibroblasts, smooth muscle cells, and peritoneal macrophages in culture. Am J Pathol 121, 200–211 (1985).

34. Romano, M. et al. An improved method on stimulated T-lymphocytes to functionally characterize novel and known LDLR mutations. J Lipid Res 52, 2095–2100 (2011).

35. Kirshner, H., Aguet, F., Sage, D. & Unser, M. 3-D PSF fitting for fluorescence microscopy: implementation and localization application. Journal of Microscopy 249, 13–25 (2013).

36. Sage, D. et al. DeconvolutionLab2: An open-source software for deconvolution microscopy. Methods 115, 28–41 (2017).

37. Vanharanta, L. et al. High-content imaging and structure-based predictions reveal functional differences between Niemann-Pick C1 variants. Traffic 21, 386–397 (2020).

38. Salo, V. T. et al. Seipin Facilitates Triglyceride Flow to Lipid Droplet and Counteracts Droplet Ripening via Endoplasmic Reticulum Contact. Developmental Cell 50, 478–493.e9 (2019).

39. Carpenter, A. E. et al. CellProfiler: image analysis software for identifying and quantifying cell phenotypes. Genome Biology 7, R100 (2006).

40. Vilhjálmsson, B. J. et al. Modeling Linkage Disequilibrium Increases Accuracy of Polygenic Risk Scores. Am. J. Hum. Genet. 97, 576–592 (2015).

41. Hunter, J. D. Matplotlib: A 2D Graphics Environment. Computing in Science Engineering 9, 90–95 (2007).

42. Michael Waskom et al. mwaskom/seaborn: v0.8.1 (September 2017). (Zenodo, 2017). doi:10.5281/zenodo.883859.

