## Supplementary methods, tables and figures for "Multiparametric Platform for Profiling Lipid Trafficking in Human Leukocytes: Application for Hypercholesterolemia"

**Detailed Methods:**

**LDL genetic risk score:** We calculated three PRSs for LDL: 1) the previously published PRS by *Talmud* et al with 12 LDL-increasing alleles, 2) a new genome-wide PRS with 6376447 variants using the recent LDpred method, and 3) a PRS s combining 1) and 2)[1,2]. The PRSs were calculated as the sum of the risk alleles weighted by their effect sizes. The weights for Talmud's PRS were based on the original publication[9].The weights for the LDpred lipid PRSs were based on a custom-run European genome-wide association study (GWAS) meta-analysis with 56945 samples excluding the FINRISK samples to eliminate sample overlap[3].The LDpred method is a Bayesian approach to calculate a posterior mean effect size for each variant based on a prior of effect size and linkage disequilibrium (a measure of how much a variant correlates with other variants)[2].Whole-genome sequences from 2690 Finns served as the linkage disequilibrium reference population for LDpred. LDpred requires a tuning parameter $\rho$ representing the fraction of causal variants in a given phenotype. We used $\rho$ of 0.01 as it provided the highest $r2$ in 4697 genotyped Finnish samples from the independent GeneRISK cohort. GeneRISK is an ongoing prospective observational study including randomly seleced 45-65 year old individuals from Southern Finland (https://thl.fi/en/web/thl-biobank/for-researchers/sample-collections/generisk-study), with the genetic risk loci based on[5]. A total of 4697 GeneRISK samples were genotyped using the HumanCoreExome BeadChip. Genotypes were called together with other available data sets using zCall at FIMM. QC and imputation were performed in the same manner as for the FINRISK samples. The PRSs were calculated using PLINK 2.0 Alpha 14. As the 12 variants included in Talmud's PRS were also included in the LDpred LDL-c PRS, we accounted for variant overlap by estimating the relative contributions of the two PRSs using linear regression with both PRSs (standardised) in a single model in the GeneRISK cohort. We combined the PRSs by weighting them by their regression coefficients and subsequently summing them together for each individual. With the combined PRS, we were not only able to account for variant overlap between the PRSs, but also address LDpred's tendency to dilute the effects of high-impact SNPs, as well as catch the non-linear contributions of the different *APOE* haplotypes to lipid levels[1,2]. We used the combined PRS in all subsequent analyses. A comparison of the different PRSs and their performance in the entire FINRISK cohort is described in Supplementary Table 3.

**Supplementary References**


1. Talmud, P. J. *et al.* Use of low-density lipoprotein cholesterol gene score to distinguish patients with polygenic and monogenic familial hypercholesterolaemia: a case-control study. *The Lancet* **381**, 1293–1301 (2013).
2. Vilhjálmsson, B. J. *et al.* Modeling Linkage Disequilibrium Increases Accuracy of Polygenic Risk Scores. *Am. J. Hum. Genet.* **97**, 576–592 (2015).
3. Surakka, I. *et al.* The impact of low-frequency and rare variants on lipid levels. *Nature Genetics* **47**, 589–597 (2015).
4. Chang, C. C. *et al.* Second-generation PLINK: rising to the challenge of larger and richer datasets. *Gigascience* **4**, (2015).
5. The CARDIoGRAMplusC4D Consortium, P. Deloukas *et al.* Large-scale association analysis identifies new risk loci for coronary artery disease. *Nature Genetics* **45**, 25–33 (2013).

**Supplementary Table 1:** Heterozygous FH patient characteristics.

| Nucleotide change | Effect on protein | LDL-c mmol/l | Cholesterol-lowering medication | Age, years | BMI, kg/m2 |
|---|---|---|---|---|---|
| c.925_931del | p.(Pro309Lysfs*59) | 2.01 | Lipcut 20 mg, Ezetrol 10 mg | 76 | 22.5 |
| c.1876G>A | p.(Glu626Lys) | 3.80 | | 74 | 26.1 |
| c.1784G>A | p.(Arg595Gln) | 2.28 | | 64 | 26.9 |
| c.1876G>A | p.(Glu626Lys) | 2.85 | Lipitor 20 mg | 63 | 23.8 |
| c.1721G>T | p.(Arg574Leu) | 3.47 | | 61 | 23.8 |
| c.925_931del | p.(Pro309Lysfs*59) | 4.77 | | 60 | 26.9 |
| c.2375T>C | p.(Ile792Thr) | 2.98 | | 66 | 23.4 |
| c.925_931del | p.(Pro309Lysfs*59) | 5.59 | Atorvastatin 80 mg Ezetrol 10 mg | 73 | 26.2 |
| c.925_931del | p.(Pro309Lysfs*59) | 4.88 | Simvastatin 20 mg | 66 | 26.6 |
| c.1085A>C | p.(Asp362Ala) | 1.89 | Crestor 10 mg | 69 | 34.0 |
| c.925_931del | p.(Pro309Lysfs*59) | 3.72 | Atorvastatin 40 mg | 70 | 23.2 |
| c.607C>T | p.(His203Tyr) | 2.49 | Lipcut 40 mg | 77 | 33.8 |
| c.1784G>A | p.(Arg595Gln) | 1.83 | THRIVE trial | 73 | 28.6 |
| c.1876G>A | p.(Glu626Lys) | 2.19 | | 61 | 22.7 |
| c.1784G>A | p.(Arg595Gln) | 3.41 | Rosuvastatin 40 mg | 58 | 28.6 |
| c.796G>A | p.(Asp266Asn) | 3.31 | Atorvastatin 40 mg | 59 | 24.7 |
| c.1876G>A | p.(Glu626Lys) | 1.73 | Atorvastatin 40 mg | 74 | 28.6 |
| c.925_931del | p.(Pro309Lysfs*59) | 3.84 | Atorvastatin 80 mg | 59 | 26.4 |
| c.1876G>A | p.(Glu626Lys) | 2.14 | Lipcut 20 mg | 67 | 25.6 |
| c.925_931del | p.(Pro309Lysfs*59) | 2.80 | Rosuvastatin 40 mg | 71 | 27.2 |
| c.1335C>A | p.(Asp445Glu) | 4.10 | | 58 | 29.0 |
| c.974G>A | p.Cys325Tyr | 6.76 | | 44 | 23.1 |
| c1739C>T | p.Ser580Phe | 5.21 | | 29 | 19.0 |

**Supplementary Table 2:** Characteristics for FINRISK subject groups with normal LDL-c (nLDL-c) and elevated LDL-c (hLDL-c) mean ± standard deviation. Hip-circ. = Hip circumference, TC = total cholesterol, LDL-c = LDL-cholesterol, HDL-c = HDL-cholesterol, TG = triglycerides, Apo-A1 = Apolipoprotein-A1, Apo-B = apolipoprotein-B. P-values were calculated with Welch's t-test.

| | **Age** year | **BMI** $kg/m^2$ | **Hip-circ.** cm | **TC** mmol/l | **LDL-c** mmol/l | **HDL-c** mmol/l | **TG** mmol/l | **Apo-A1** mmol/l | **Apo-B** mmol/l |
|---|---|---|---|---|---|---|---|---|---|
| **nLDL-c** n = 19 | 58.8 ±10.0 | 25.3 ±4.3 | 99.0±10.3 | 4.5 ±0.7 | 2.3 ± | 1.7 ±0.1 | 1.3 ±1.5 | 1.7 ±0.4 | 0.6 ±0.1 |
| **hLDL-c** n = 20 | 61.3 ±13.4 | 27.1 ±2.6 | 101.3 ±7.7 | 8.1 ±1.2 | 5.8±1.1 | 1.6 ±0.4 | 1.6 ±0.5 | 1.6 ±0.3 | 1.5 ±0.3 |
| p-value | n.s. | n.s. | n.s. | <0.001 | <0.001 | n.s. | n.s. | n.s. | <0.001 |

**Supplementary Table 3. Performance of PRSs for LDL-c in the entire FINRISK cohort:**

Comparison of the performance of Talmud's 12-SNP PRS[1], the LDpred PRS, and the PRS combining Talmud's and LDpred PRSs for LDL-C in the FINRISK cohort with lipid measurements. Performances were estimated using linear regression with residual lipid measurements after adjusting for age and sex as the response. Weight in combined PRS refers to the regression coefficients of standardised Talmud's and LDpred LDL-C PRSs estimated in the independent GeneRISK cohort (n = 4697). PRS, polygenic risk score. SE, standard error. LDL-C, LDL-cholesterol. TG, triglycerides. AIC, Akaike information criterion.

| | ***β*** | **SE** | ***p*** | **Adjusted $r^2$** | **AIC** | **Weight in combined PRS** |
|---|---|---|---|---|---|---|
| **LDL-C** | | | | | | |
| Talmud's PRS | 0.24 | 0.0060 | < 0.0001 | 0.061 | 65102 | 0.084 |
| LDpred PRS | 0.29 | 0.0059 | < 0.0001 | 0.091 | 64683 | 0.244 |
| Combined PRS | 0.30 | 0.0059 | < 0.0001 | 0.098 | 64126 | |

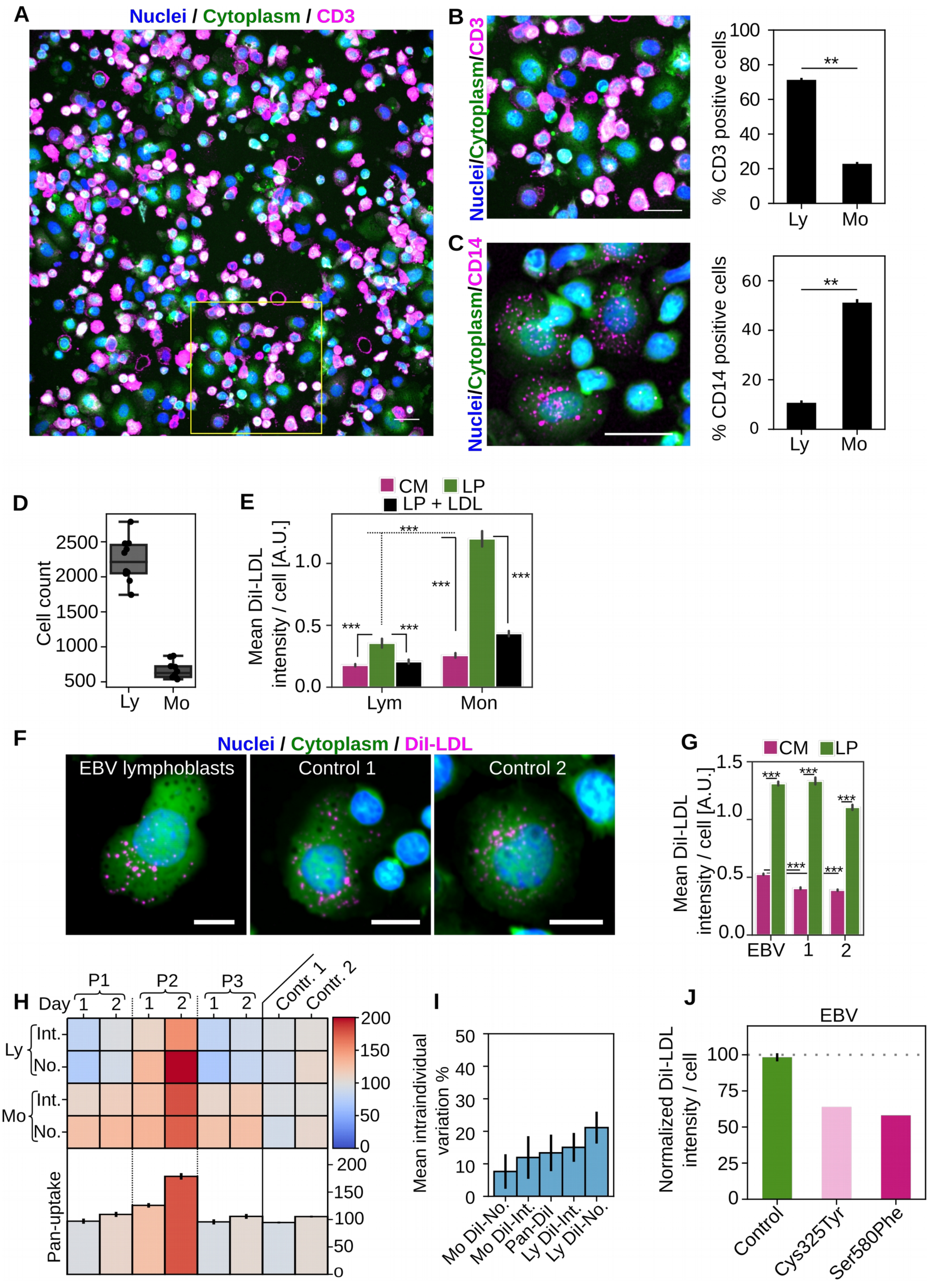
A
Nuclei / Cytoplasm / CD3
B
Nuclei/Cytoplasm/CD3
% CD3 positive cells
Ly
Mo
C
Nuclei/Cytoplasm/CD14
% CD14 positive cells
D
Cell count
E
CM
LP
LP + LDL
Mean DiI-LDL intensity / cell [A.U.]
Lym
Mon
F
Nuclei / Cytoplasm / DiI-LDL
EBV lymphoblasts
Control 1
Control 2
G
Mean DiI-LDL intensity / cell [A.U.]
EBV
H
P1
P2
P3
Contr. 1
Contr. 2
Day
Int.
No.
Pan-uptake
I
Mean intraindividual variation %
Mo DiI-No.
Mo DiI-Int.
Pan-DiI
Ly DiI-Int.
Ly DiI-No.
J
EBV
Normalized DiI-LDL intensity / cell
Control
Cys325Tyr
Ser580Phe

**Supplementary Figure 1 A**) Representative image of PBMC cells stained with DAPI (nuclei), CellMask Green (cytoplasm) and anti-CD3 antibodies (lymphocytes). **B**) Zoom in view of the yellow area in (**a**) and automated quantification of % CD3 positive cells for cells with a cytoplasm area below 115 $\mu m^2$ (designated as lymphocyte population) and above 115 $\mu m^2$ (designated as monocyte population), n = 2 controls, each containing PBMCs from 4 individuals; ±SEM. **c**) Representative zoom in image of anti-CD14 stained PBMCs and quantification of % CD14 positive cells in lymphocyte (Ly) and monocyte (Mo) populations as defined in (**B**), n = 2 controls, each containing PBMCs from 4 individuals; ±SEM. **D**) Box plot for lymphocyte and monocyte cell counts per well for a control containing PBMCs from 4 individuals; representative of eight independent experiments, n = 8 wells. **E**) Quantification of DiI-LDL intensities for lymphocyte and monocyte populations after treatment with control medium (CM, 10%FBS), lipid starvation (LP) or lipid starvation medium supplemented with 100 µg / ml native LDL during DiI-LDL uptake phase (LP+LDL). On average 5240 lymphocytes and 2580 monocytes were analyzed for each treatment per sample; ± 95%CI. **F)** DiI-LDL uptake in control EBV lymphoblasts (**EBV**) and PBMCs of two controls after lipid starvation. Representative images are shown. **G**) Quantification of mean DiI-LDL intensities in EBV lymphoblasts and monocytes from two controls after treatment with control medium (CM) or lipid starvation medium (LP), n > 20 000 for EBV lymphoblasts, and >16 000 monocytes from 10 independent experiments; ± 95% CI. **H**) Lymphocyte and monocyte cellular DiI-LDL intensities (Int), DiI-LDL organelle counts (No), and the pan-uptake score for three individuals, sampled on two consecutive days; ±SEM, n = 8 wells (32 for pan-uptake) measured in two independent measurements. **I**) Average intraindividual variation for monocyte and lymphocyte uptake scores and pan-uptake, n = 3 individual persons; ±SEM. **J**) Quantification of cellular DiI-LDL intensities in EBV lymphoblasts from a control and two familial hypercholesterolemia patients (FH) after 72 h of lipid starvation; ±95% CI.

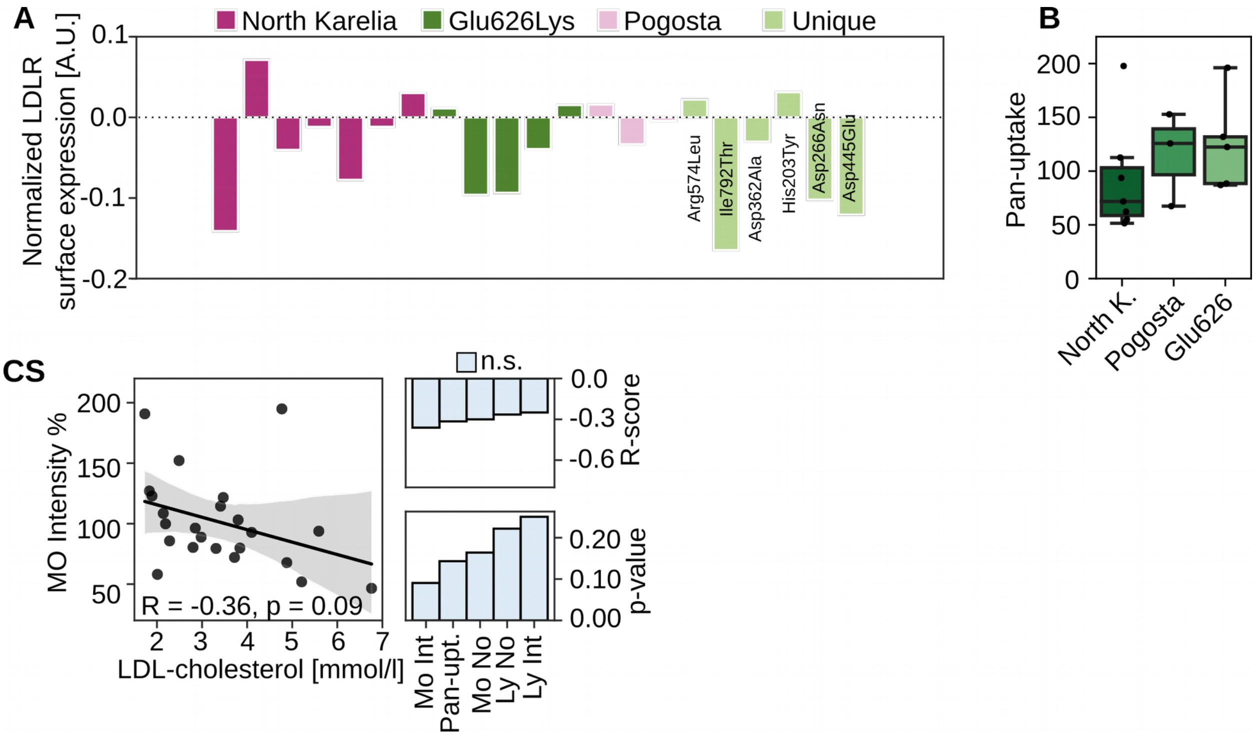


**Supplementary Figure 2 A)** Quantification of LDLR surface expression in monocytes after 24 h lipid starvation, relative to controls. On average, 1175 monocytes were quantified for each patient. **B**) Box plots for pan-uptake in heterozygous FH variant groups, North Karelia (North K.)(n = 7), Pogosta (n = 3) and Glu626 (n = 5). **C**) Correlation of monocyte DiI-LDL intensity and LDL-c concentration for heterozygous FH patients (n = 23) together with R- and p-values for all LDL uptake scores. Grey areas in scatter plots indicate 95% CI.

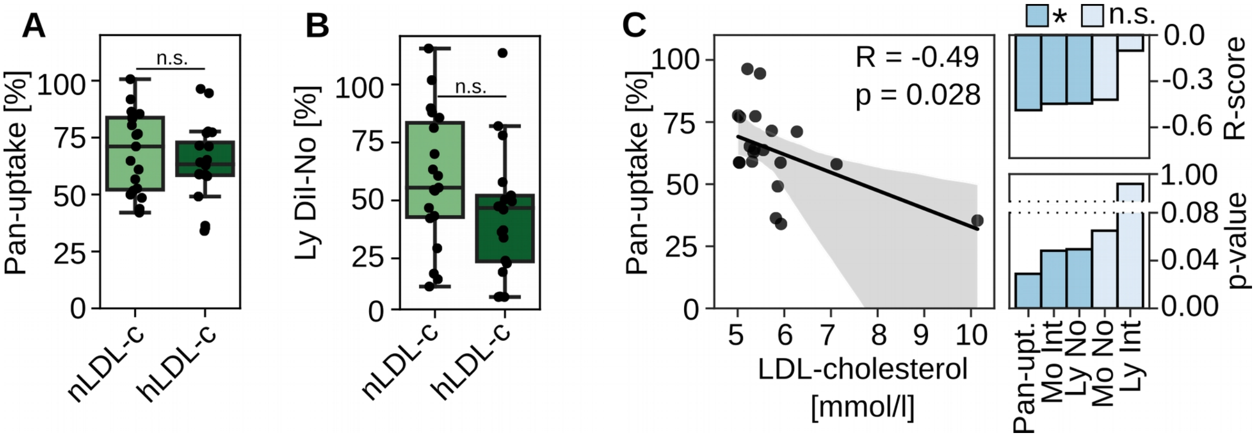


**Supplementary Figure 3)** Box plots for pan-uptake (**A**) and lymphoyte (Ly) DiI-LDL organelle numbers (DiI-No) (**B**) in individuals with normal (nLDL-c, LDL-c 2-2.5 mmol/l) and elevated LDL-cholesterol (hLDL-c, >5 mmol/l LDL-c); nLDL-c, n =19; hLDL-c, n =20, Welch's t-test. **C**) Correlation of pan-uptake with LDL-cholesterol for hLDL-c subjects, including R- and p-values for individual LDL uptake scores; n = 20. Grey areas in scatter plots indicate 95% CI.

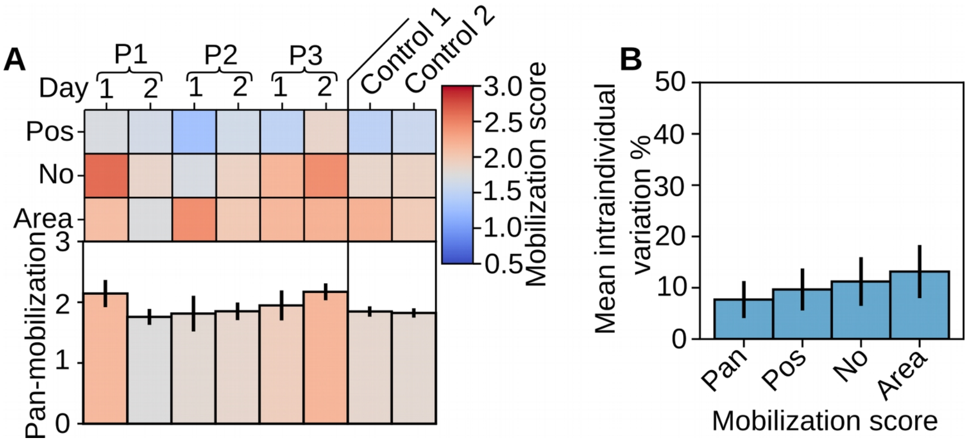


**Supplementary Figure 4)** Intraindividual variation of lipid mobilization scores. **A)** Quantification of monocyte lipid mobilization scores LD-Pos, LD-No, LD-Area- and pan-mobilization as described in (**Figure 4G**) for three individuals sampled on two consecutive days; n = 4 wells (12 wells for pan-mobilization) from two independent measurements. **B**) Average intraindividual variation for lipid mobilization scores, n = 3 individual persons; ±SEM.

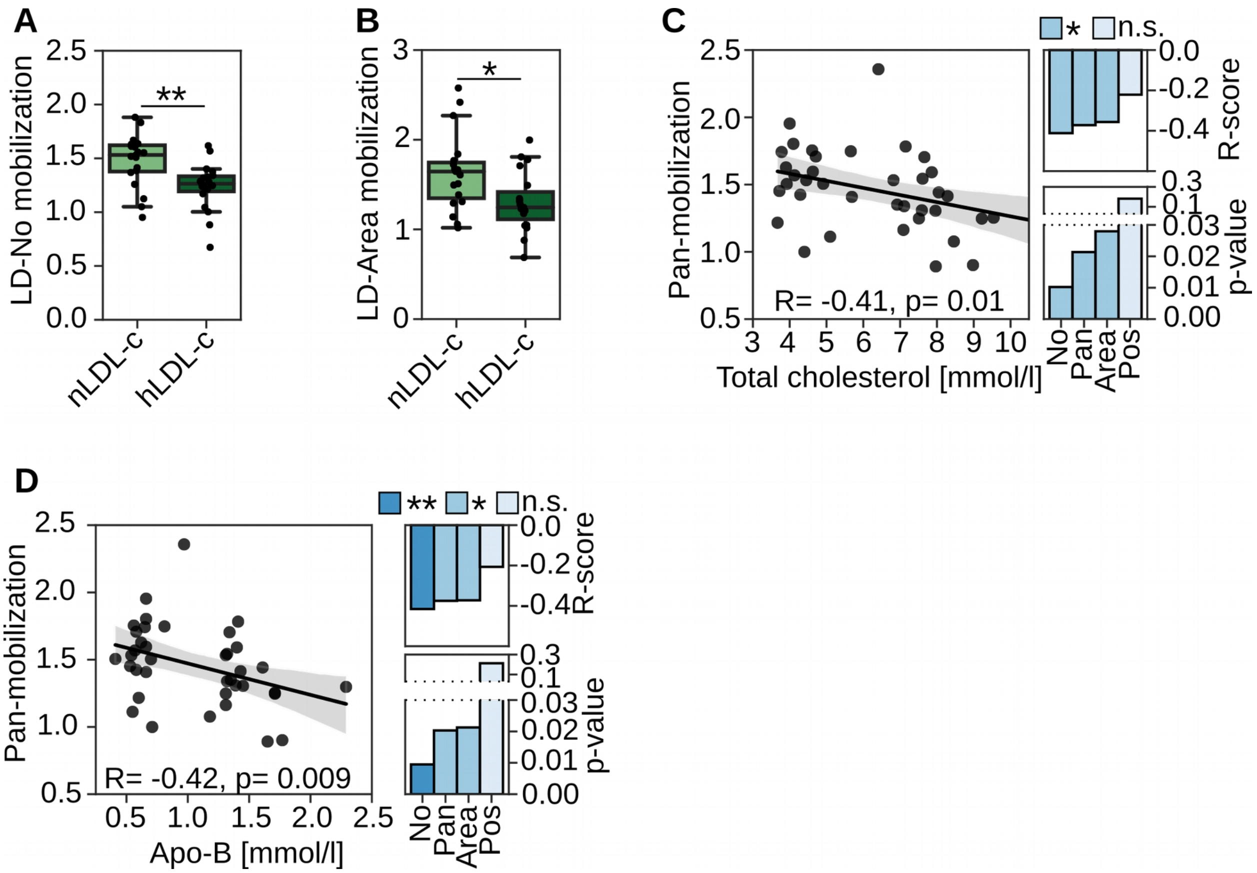


**Supplementary Figure 5)** Box plots for lipid mobilization scores LD-No **(A)** and LD-Area **(B)** in individuals with normal (nLDL-c, LDL-c 2-2.5 mmol/l) and elevated LDL-cholesterol (hLDL-c, >5 mmol/l LDL-c); nLDL-c, n =19; hLDL-c, n =19; Student's t-test. ** p<0.01, **p<0.05. **C**) Correlation of pan-mobilization with total-cholesterol (mmol/l) and Apolipoprotein-B, Apo-B (mmol/l) (**D**), including R- and p-values for all mobilization scores. Grey areas in scatter plots indicate 95% CI.

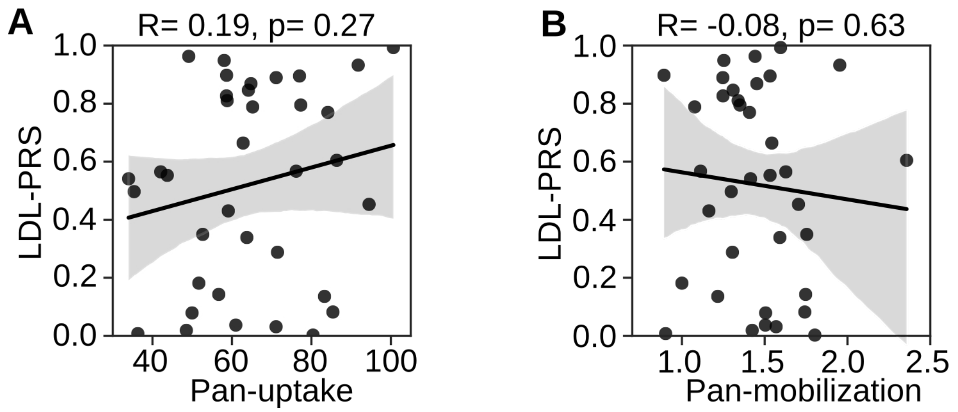


**Supplementary Figure 6)** A polygenic risk score for high LDL-cholesterol (LDL-PRS) does not correlate with pan-uptake (**A**) and pan-mobilization (**B**) scores: n = 37. Grey areas in scatter plots indicate 95% CI.
